# Characterizing RNA-binding Ligands on Structures, Chemical Information, Binding Affinity and Drug-likeness

**DOI:** 10.1101/2022.01.03.474753

**Authors:** Cong Fan, Xin Wang, Tianze Ling, Huiying Zhao

**Author notes:** **Contact information of corresponding authors:** Huiying Zhao, PhD, Department of Medical Research Center, Sun Yat-Sen Memorial Hospital, 107 Yan Jiang West Road Guangzhou P.R. China, 500001; Fax: 510220.

## Abstract

Recent studies suggest RNAs playing essential roles in many cell activities and act as promising drug targets. However, limited development has been achieved in detecting RNA-ligand interactions. To guide the discovery of RNA-binding ligands, it is necessary to characterize them comprehensively. We established a database, RNALID that collects RNA-ligand interactions validated by low-throughput experiment. RNALID contains 358 RNA-ligand interactions. Comparing to other databases, 94.5% of ligands in RNALID are completely or partially novel collections, and 51.78% have novel two-dimensional (2D) structures. The ligand structure analysis indicated that multivalent ligands (MV), ligands binding with cellular mRNA (mRNA), ligands binding with RNA from virus (vRNA) and ligands binding with RNA containing repetitive sequence (rep RNA) are more structurally conserved in both 2D and 3D structures than other ligand types. Binding affinity analysis revealed that interactions between ligands and rep RNA were significantly stronger (two-tailed MW-U test P-value = 0.012) than the interactions between ligands and non-rep RNAs; the interactions between ligands and vRNA were significantly stronger (two-tailed MW-U test P-value = 0.012) than those between ligands and mRNA. Drug-likeness analysis indicated that small molecule (SM) ligands binding with non-rep RNA or vRNA may have higher probability to be drugs than other types of ligands. Comparing ligands in RNALID to FDA-approved drugs and ligands without bioactivity indicated that RNA-binding ligands are different from them in chemical properties, structural properties and drug-likeness. Thus, characterizing the RNA-ligand interactions in RNALID in multiple respects provides new insights into discovering and designing druggable ligands binding with RNA.

**KEY POINTS:**

- We established a database, RNALID, by collecting RNA-ligand interactions validated by low-throughput experiments, which contains 358 RNA-ligand interactions, and 94.5% of them are completely or partially novel collections.
- Ligands in RNALID were divided into fourteen types, and were analyzed on chemical properties, spatial properties, binding affinity and drug-likeness. Small molecule (SM) ligands binding with non-repeat RNA or virus RNA were indicated having higher probability to be drugs than other types of ligands.
- Ligands in RNALID were different from protein-binding ligands and ligands without bioactivity in chemical properties, structural properties and drug-likeness.

## INTRODUCTION

Protein-ligands interactions discovered in traditional paradigm potentially fail to meet the urgent disease-therapy needs because protein-coding genes take up only ∼1.5% of human genome and only 10%-15% of them are druggable^1^. Comparably, human genome produces a huge number of non-coding RNAs, for example, probably more than 15,000 long non-coding RNAs^2^, and a subset of these may include many good drug targets. Ligands targeting on RNAs may be more likely to involve in cellular regulation than ligands targeting on proteins^3,4^. Detecting ligands targeting on RNAs provides a promising approach for drug discovery^5^. Recent studies presented small molecules and oligonucleotides for cancer treatment in clinical trials by modulating the mRNA spliceosome^6-8^. Other studies have reported the roles of microRNA and long non-coding RNA (lncRNA) in human diseases ^9-13^. By interacting with these non-coding RNAs, small molecules have been discovered as playing functions in anti-diseases^13, 14^. Until now, only six ligands (Azacitidine, Carmustine, Fluorouracil, Capecitabine, Tegafur-uracil and Cocarboxylase) interacting with RNAs have been approved as drugs by FDA (https://www.drugbank.ca/). Thus, it is necessary to identify more RNA-targeting drugs.

A plenty of studies have been performed to identify RNA-ligand interactions experimentally. For instance, Serganov et.al presented the crystal structures of the Fusobacterium nucleatum riboswitch bound to riboflavin and antibiotic roseoflavin^15^. However, there are few databases integrating these published RNA-ligand interactions. NALDB^16^ is a database including experimentally validated interactions between small molecules and various types of nucleic acids. In NALDB, 104 entries of RNA-ligand interactions were included. However, only information of ligands can be accessed but not RNAs. R-BIND^17-19^ is a more recently developed database containing information on reported chemical probes that target on non-ribosomal RNAs. Totally, 116 bioactive ligands were included in R-BIND. Nevertheless, R-BIND lacks several aspects of information on ligands. Firstly, in R-BIND, only chemical information but not ligand structural information was available, whereas studies have revealed that ligands binding with the same RNA are more likely to share similar two-dimensional structures^20^. Secondly, R-BIND divided ligands simply into small molecules (SM) and multivalent ligands (MV), and ignored the difference of ligands in other repects^18^. Moreover, sequences of RNA binding with ligands as well as the therapeutical destinations of the ligands were not included in R-BIND database. Using the data of R-BIND, *Warner et*.*al*. analyzed RNA-ligand interactions on relationships between drug-like scores and binding affinities^5^. This analysis is based on a small sample of totally 30 ligands because of the prerequisite that each sample ligand should bind to RNAs with known superior structures. Currently, no studies have characterized the RNA-ligand interactions on their 2-dimensional (2D) and 3-dimensional (3D) structures, chemical properties and spatial properties by dividing ligands into various types. Furthermore, no studies have been performed to reveal the relationship between RNA-ligand binding affinities and drug-likeness of ligands.

In this study, we manually curated a database, RNALID, including experimentally validated RNA-ligand interactions published before Feb, 2020. The database contains totally 358 experimentally validated RNA-ligand interactions, in which 340 entries are partially new (added new information for previously collected RNA-ligand interactions) or completely new (newly collected RNA-ligand interactions) comparing to existing databases. A webservice was provided to predict ligand-RNA interactions through comparing the query on ligand structures, chemical properties, RNA sequence and RNA secondary structure to the ligands or RNAs in RNALID. Using the RNALID, comprehensive analysis was performed on 2D and 3D structural characteristics, spatial properties and chemical properties of RNA-binding ligands. Moreover, this study revealed the relationships between RNA-ligand binding affinities and 28 comprehensive drug-like indexes. To identify the differences between RNA-binding and non-RNA-binding ligands, we collected 42,234 ligands without bioactivates and 7,753 FDA approved non-RNA-binding drugs, and compared them with the RNA-binding ligands in RNALID on chemical properties, spatial properties, structures and drug-likeness.

## MATERIAL AND METHODS

### Collection of RNA-ligand interactions

RNA-ligand interactions were collected in RNALID if the interactions were validated by low-throughput experiments according to published studies in PubMed. The low-throughput experiments include Isothermal Titration Calorimetry (ITC), spectroscopic deciphering, circular dichroism (CD) spectroscopy, Fluorescence-based binding assay, NMR, gel mobility shift assays, CD spectroscopy, and UV spectroscopy. Interactions involving only rRNAs were ignored^17, 19^. We searched PubMed with keywords “RNA”, “drug”, “ligand” and “interaction” along with their plural forms. Totally, 103 literatures were involved in this study. From these literatures, we manually extracted information on ligand structures, sequences of RNA binding with ligands, HGNC names of binding RNAs, and targeted diseases of ligands. If multiple ligand analogs appeared in one publication, the ligand with representative structure or highlighted more than once by the author in the abstract or discussion section would be adopted. The graphical illustrations of ligand structure were converted to SMILES format by BKChem (http://bkchem.zirael.org/). If a PDB ID (http://www.rcsb.org/) of the ligand was offered by the author, the SMILES and InChIKey on the PDB webpage were both kept in RNALID. If the ligand has been included in the PubChem database (https://pubchem.ncbi.nlm.nih.gov/), the SMILES and InChIKey of it in the PubChem were kept.

To integrate RNA-ligand interactions in RNALID with other databases, we manually curated RNA sequence, HGNC names of corresponding RNA and targeted diseases for RNA-ligand interactions in R-Bind, the only existing database containing downloadable RNA-ligand interactions. All ligands in R-BIND were transformed from SMILES format to InChIKeys using OpenBabel software^21^.

### Ligand classification

Ligands in RNALID were grouped under a variety of classification principles as shown in Table S1. In detail, using the principle of structural characters, ligands were defined as “MV” (multivalent ligands) if their structures were in the presence of an alkyl, aryl, or peptide-like linker between two or more binding moieties. Otherwise, they were defined as “SM” (small molecules). Using the principle of target preference, ligands in format of InChIKeys in the BindingDB database^22, 23^ (https://www.bindingdb.org/bind/index.jsp) were considered as binding with both RNAs and proteins (“RNA&protein” type) while the remained ligands were considered as only binding with RNAs (“RNA” type). Ligands were classified as “rep RNA” if they bound with at least one RNA with repetitive sequence, otherwise they were defined as “non-rep RNA” type. Finally, according to RNA types binding with them, ligands were classified into six groups including mRNA, vRNA, miRNA, tRNA, ncRNA, siRNA, and others. Here, vRNA represents mRNA from virus, and mRNA represents mRNA from cellular. To obtain statistically meaningful results, the analysis focused on ligands binding with mRNA and vRNA because their sample sizes are more than 20 (Table S1.).

### Ligand similarity

In this study, ligand similarities include chemical similarity and structural similarity. To evaluate the chemical similarity between ligands, 12 cheminformatic parameters for each ligand were generated by RDKit (http://www.rdkit.org/) (Table S2). Chemical similarities between ligands based on these cheminformatic parameters were calculated by procedures shown in Supplementary material.

The ligand structure similarities were evaluated by 2D structure similarly and 3D structure similarity, respectively. The calculation of 2D structure similarity was performed by the procedure described in a previous study^24^, where ligand structures were firstly transformed to fingerprint format of FP2 by OpenBabel^21^. The 3D structure similarity of ligands was evaluated by LSalign^25^ under the mode of flexible alignment as described in a previous study^26^.

### Spatial property of ligand

The spatial property of ligand was evaluated by the principal moment of interias (PMI). The PMI was performed using a functional module in RDKit. The detailed steps are included in the Supplementary material.

### Drug-like index

Totally, 28 drug-like indexes were considered in this study (Table S3). They were calculated by CMPNN algorithm^27^ with training data from databases of BBBP (http://moleculenet.ai/datasets-1), Tox21 (https://tox21.gov/), Sider (http://sideeffects.embl.de/), ClinTox (https://www.clintox.org/), ESOL^28^ and FreeSolv^29^. The differences of drug-like index scores between ligands types were evaluated by Mann-Whitney U test.

### Collection of FDA drugs and ligands without bioactivities

FDA drugs were collected from the DrugBank database^30-34^, which contains drugs approved by Food and Drug Administration (FDA). They were filtered by Lipinski’s rule of five (RO5)^35^ to obtain the protein-binding drugs^5^. Ligands without bioactivates were collected from the Bioassay&Bioactivities section of PubChem project (https://pubchem.ncbi.nlm.nih.gov/). These ligands were named as “control ligand” in this study.

## RESULTS

### Establishing an RNA-ligand interaction database (RNALID)

RNALID database (http://121.46.19.89/RNALID/html/index.html#/database) contains 358 experimentally validated RNA-ligand interactions, in which 74.86% of RNAs have sequence information, 11.73% of ligands have drug therapy information obtained from DRUGBANK ^30-34^ (Table S4), 30.17% of RNAs have HGNC symbols, 41.9% of RNAs or ligands bind with other types of targets except ligands or RNAs, and 93.58% of RNA-ligand interactions have information on associated diseases (Table S5 and Figure S2). All this information is not available yet in existing databases. Additionally, RNALID includes binding affinity for 58.66% of RNA-ligand interactions (in at least one record of K_i_, K_d_, IC_50_, EC_50_ and CC_50_). The chemical and structural properties of ligands in RNALID were compared with other available RNA-ligand interactions databases as shown in Supplementary material, identifying 51.78% of ligands have novel 2D structures comparing to other databases when using 2D structure similarity score > 0.4 to define ligands with similar structures (Figure S4). The description of user interface of RNALID is in Supplementary material. Ligands in RNALID were divided into 14 types as shown in Table S1.

### Chemical properties of different types of ligands in RNALID

The varieties in cheminformatic parameters (Table S2) of different types of ligands were evaluated by MW-U test. The cheminformatic parameters of SM and MV ligands were compared in Figure 1A. Totally, 9 out of 12 cheminformatic parameters, including “Molecular weight”, “tPSA”, “ArRings”, “HeRings”, “HBA”, “HBD”, “Rings”, “RotB”, and “ASA” are significantly different in SM and MV ligands. MV ligands show significantly higher values than SM ligands in all these 9 cheminformatic parameters, which indicates MV ligands have higher molecular weight, topological polar surface area (tPSA), Rings, aromatic rings (ArRings), heteroatom-containing rings (HetRings), rotatable bonds (RotB), ASA, as well as hydrogen-bond acceptors (HBA) and donors (HBD). According to Lipinski’s rule of five (RO5)^35^, orally bioavailable ligands have molecular mass less than 500Da (MWT ⩽ 500), fewer hydrogen-bond donors and hydrogen-bond acceptors (H-bond donors ⩽ 5, H-bond acceptors ⩽ 10), and less octanol-water partition coefficient (log P ⩽ 5)^35^. Thus, MV ligands may have lower probability to be oral bioavailable ligands than SM ligands.

**Figure 1.**
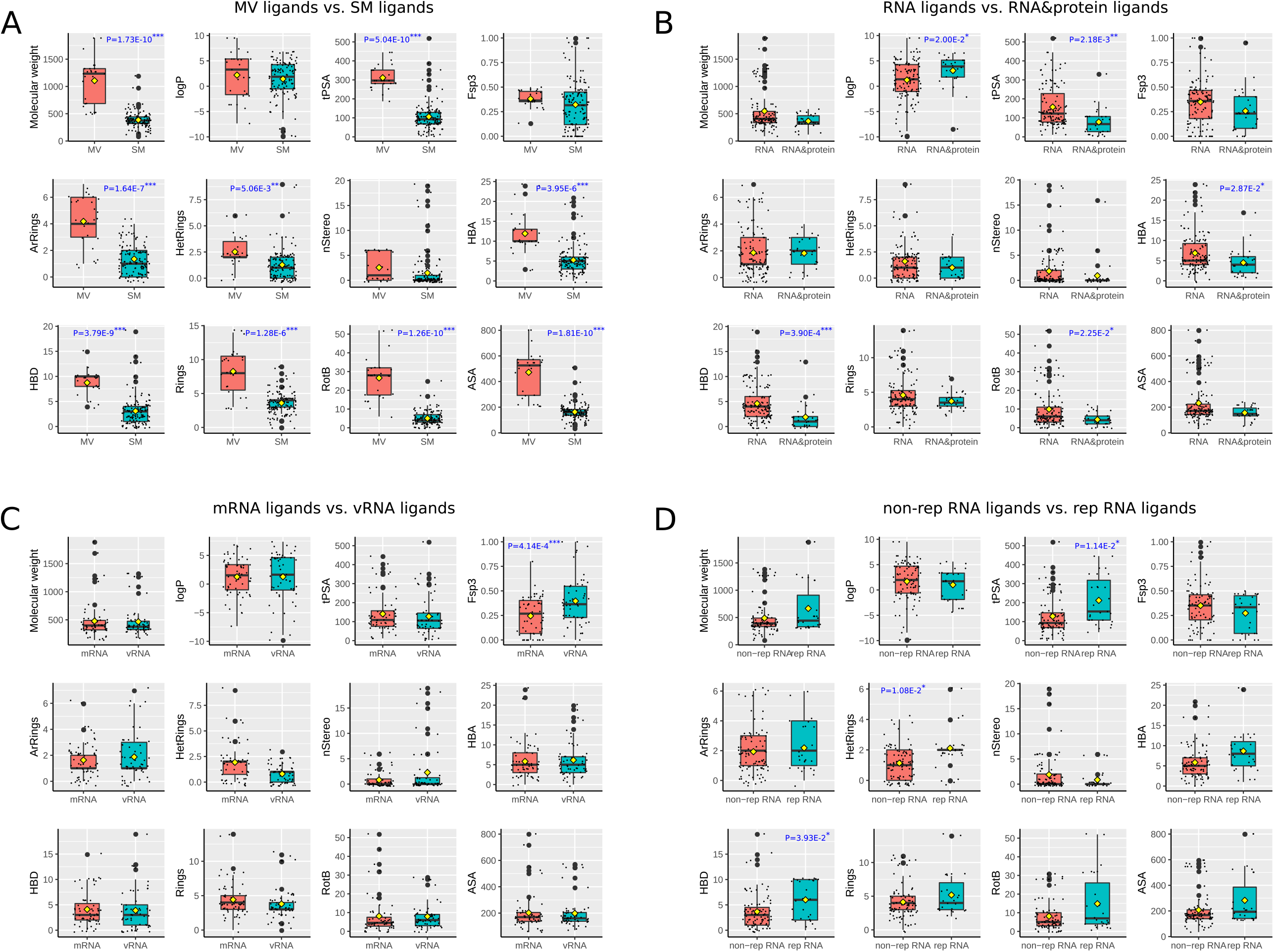
Comparing chemical characters of ligands in RNALID. **(A)** Comparing SM ligands and MV ligands. **(B)** Comparing ligands only binding with RNA, and ligands binding with both RNA and protein. **(C)** Comparing ligands binding with mRNA from cellular and ligands binding with mRNA from virus (vRNA). **(D)** Comparing ligands binding with RNA including repeat sequence (rep RNA) and ligands binding with RNA without repeat sequence (non-rep RNA). The differences of chemical characters of ligands were evaluated by MW-U test. “*” denotes P-value < 0.05, “**” denotes P-value < 0.01, “***” denotes P-value < 0.001.

Comparing ligands only binding with RNA (RNA type) and ligands binding with both RNA and protein (RNA&protein type) on cheminformatic parameters, we found the RNA&protein type of ligands exhibited a significant increasing in logP while a reduction in polar surface area (tPSA), hydrogen bond donors (HBD), rotatable bonds (RotB), and hydrogen-bond acceptors (HBA) than ligands only binding with RNA (Figure 1B). This result indicated that ligands binding with both RNA and protein have a higher possibility to be oral bioavailable ligands according to rules of RO5^35^ than ligands only binding with RNA.

Ligands interacting with mRNA have significantly lower hybridized carbons (Fsp^3^) values than ligands binding with vRNA while they show similar values in the other 11 cheminformatic parameters (Figure 1C).

Comparing ligands binding with rep RNA to ligands binding with non-rep RNA indicated that ligands binding with rep RNAs have significantly (P-value = 1.14E-2) higher polar surface area (tPSA), a greater number of HetRings (P-value = 1.08E-2) and HBD (P-value = 3.93E-2) than ligands binding with non-rep RNAs (Figure 1D).

### Structure diversity of different types of ligands

In order to evaluate the structure diversity of ligands, we calculated pair-wised 2D structure and 3D structure similarities of all ligands in RNALID. The pair-wised structure similarities for the same type of ligands were defined as intra-similarity. We calculated the average scores of the intra-similarity. The results were shown in Figure 2. MV ligands have the highest intra-similarity in 2D structures than other types of ligands. The average 2D structure similarity score of MV ligands was significantly (P-value = 7.61E-24) higher than that of SM ligands. Thus, ligands in MV group may be less diverse in 2D structures than ligands in other types. The average 3D structure similarity score of ligands binding both RNA and protein was significantly (P-value = 1.63E-4) higher than that of ligands binding only with RNAs, indicating the ligands binding with both RNA and protein are less structurally diverse than ligands only binding with RNAs. Moreover, ligands binding with non-rep RNA have significantly (P-value = 4.69E-11 and 6.43E-4) lower 2D and 3D structure similarity than the ligands binding with rep RNA, suggesting a higher structural conservation of ligands binding with repeated RNA sequence.

**Figure 2.**
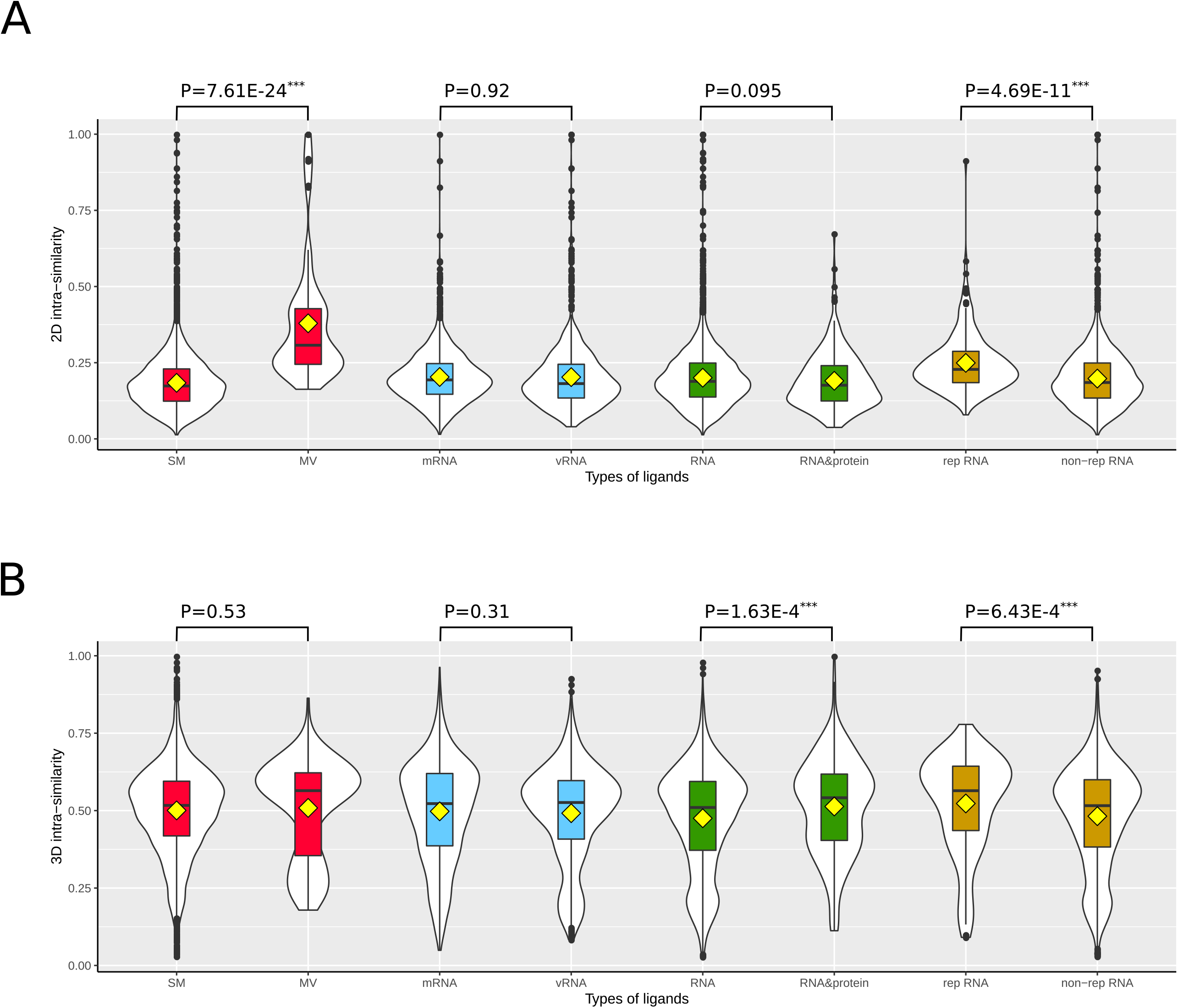
Comparing 2D and 3D structure similarity of ligands in RNALID. **(A)** Comparing 2D structures of the same type of ligands. **(B)** Comparing 3D structures of the same type of ligands.

We further calculated the pair-wised structure similarities of ligands across different types. These structure similarity scores were defined as inter-similarity. The results were shown in Figure S5. We found that ligands in types of MV, mRNA, vRNA and rep RNA exhibited higher intra-similarity scores than inter-similarity scores in both 2D and 3D structures, suggesting they are more structurally conserved.

### PMI analysis on ligands in different groups

Spatial properties of ligands were evaluated by PMI analysis. As shown in Figure 3, ligands binding with mRNA from cellular are mainly near rod vertex while ligands binding with vRNA are close to disc vertex (Figure 3A). However, the cumulative fraction graphs (Figure S6A) did not confirm the statistical significance between these two types of ligands. When ligands were grouped into ligands solely binding with RNA and binding with both RNA and protein (RNA&protein), they were found sharing the same spatial properties and mainly located near rod and disc vertex (Figure 3B). These conclusions were verified by cumulative fraction graphs. Meanwhile, neither of them preferred sphere-like shape (Figure S6B). The analysis was finally performed for the ligands belonging to “rep RNA” and “non-rep RNA” groups, respectively. Ligands binding with rep RNA were largely populated in the region near rod vertex while the ligands binding with non-rep RNA group were mainly located near disc vertex (Figure 3C). Meanwhile, the cumulative fraction graphs indicated that ligands binding with rep RNA were more rod-like ligands while ligands binding with non-rep RNA were more disc-like ligands (Figure S6C).

**Figure 3.**
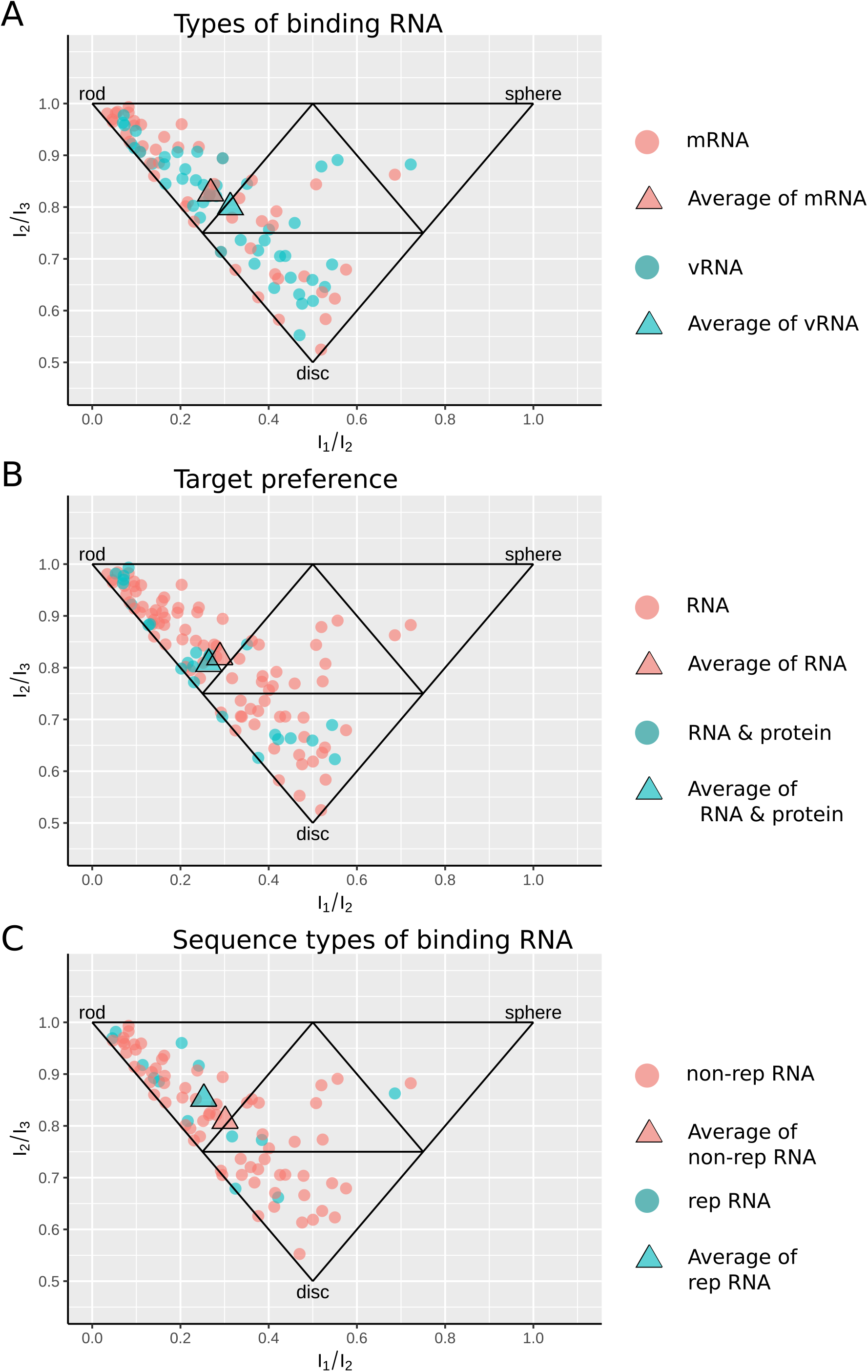
Comparing spatial properties of different types of ligands in RNALID. **(A)** Comparing ligands binding with mRNA and ligands binding with vRNA. **(B)** Comparing the spatial properties of ligands only binding with RNA to the ligands binding with both RNA and protein. **(C)** Comparing ligands binding with RNA with repeat sequence and ligands binding with RNA without repeat sequence. Each point represents one ligand. A triangle (average) represents the mean values of coordinates in the 2D-space formed by I_1_/I_3_ and I_2_/I_3_for each type of individual ligands.

### Binding affinity of RNA-ligand interactions for ligands belonging to different groups

In this study, we collected binding affinities for 204 RNA-ligand interactions, which are in IC_50_, EC_50_, K_i_, K_d_ or CC_50_ format. As shown in Figure S7A, significant difference (two-tailed MW-U test P-value = 0.012) in binding affinity is observed between ligands binding with rep RNAs and ligands binding with non-rep RNAs. The interactions between ligands binding with rep RNAs were significantly stronger than the interactions between ligands binding with non-rep RNAs. Meanwhile, the interactions between ligands and vRNAs were significantly (two-tailed MW-U test P-value = 0.012) stronger than those between ligands and mRNA (Figure S7B). No significant differences were observed between MV ligands and SM ligands (Figure S7C), and between ligands binding with only RNA and ligands binding with RNA&proteins (Figure S7D).

### Relationships between drug-like indexes and RNA-ligand binding affinities

As shown in Figure 4A and Figure S8, ligands binding with mRNA from cellular have significantly higher QED scores for drug-like indexes of SR.p53, SR.MMP, SR.HSE, SR.ATAD5, SR.ARE, NR.PRAR.gamma, NR.ER.LBD, NR.ER, NR.Aromatase, NR.AR.LBD, NR.AhR and LIVER than ligands binding with coding RNA from virus (vRNA). This result indicates that vRNA ligands process higher drug-likeness than mRNA ligands because druglike index of LIVER (drug metabolism in liver) is positively related to unstable and unpredictable bio-effects of drugs while the other druglike indexes are positively related to toxicity. Ligands binding with rep RNA show significantly higher QED scores for drug-like indexes of SR.ATAD5, NR.ER, NR.AhR, and LIVER than ligands binding with non-rep RNA. Because higher QED scores of these druglike indexes were positively related with toxicity or unstable drug effects, non-rep RNA ligands exhibit higher drug-likeness. Ligands binding with RNA & proteins have significantly higher QED scores for drug-like indexes of LOGP, CYP2C9_S, CLINTOX and CACO2 than ligands solely binding with RNA. Finally, ligands in MV are indicated with higher QED scores for drug-like indexes of SR.p53, SOLUBILITY, NR.PRAR.gamma, NR.AR.LBD, CYP3A4_S and CYP2D6_I than ligands in SM. Four out of the five indexes were positively related with toxicity. Thus, SM ligands may have higher probability to be drugs than MV ligands. This is in consistent with the findings in comparing cheminformatic parameters of these two types of ligands.

**Figure 4.**
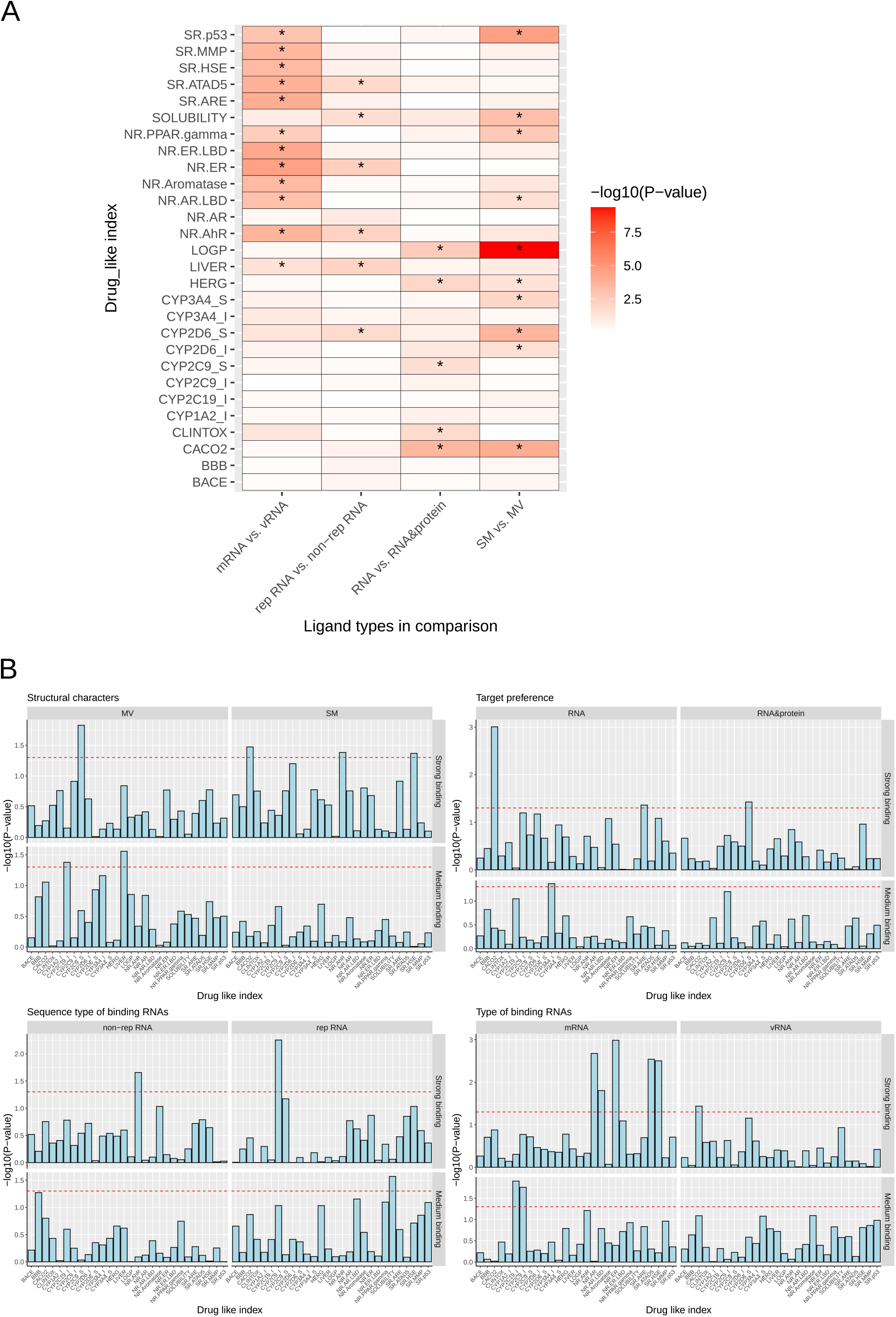
Relationships between RNA-ligand binding affinity and drug-like indexes. **(A)** Comparison of QED scores among different types of ligands (two-tailed MW-U test was used to evaluate the significance). **(B)** P-values of Pearson correlation coefficient of RNA-ligand binding affinity and QED scores for different types of ligands.

For each type of ligands, we examined the correlation between the binding affinity and QED scores of drug-like indexes (Figure 4B and Figure S9). Figure 4B shows the P-values of Pearson Correlation Coefficient (PCC) between QED score and binding affinity of ligand and RNA, and indicated that the binding affinities of strong RNA-ligand interactions [binding affinity lower than 1 *μ M* (1E-6 *mol/L*) and higher than 0 *nM*] were more likely to be significantly correlated with QED scores than the medium interactions (binding affinities were higher than 1 *μ M* and lower than 1 *mM*). The detailed definitions of strong and medium RNA-ligand interactions were shown in Supplementary Material. The QED scores of 10 drug-like indexes, CACO2, NR.ER, NR.AR, SR.ATAD5, SR.HSE, CYP2C9_S, NR.AR.LBD, NR.AhR, CYP2D6_S and SR.ARE were found significantly (P-value = 9.83E-4, 1.00E-3, 2.10E-3, 2.87E-3, 4.29E-2, 1.50E-2, 1.57E-2, 2.20E-2, 3.74E-2 and 4.34E-2, respectively.) correlated with binding affinities of strong interactions while the QED scores of two drug-like indexes, CYP3A4_S and LIVER were found significantly correlated with binding affinities of medium interactions (P-values=3.09E-2 and 2.77E-2, respectively).

When the binding affinities were strong interactions, the binding affinities of SM ligands were found significantly correlated with the QED scores of NR.AhR, CACO2, and SR.HSE. The binding affinities of MV ligands with strong interactions were found correlated with the QED score of CYP2C9_S only. In comparison, the binding affinity of MV ligands with medium interactions were found significantly correlated with the QED scores of LIVER while the binding affinity of SM ligands with medium interactions were not found significantly correlated with any QED scores. Thus, the binding affinities of MV and SM ligands were possibly influenced by different drug-like index. Figure S9 shows the scatter plots describing the correlations between binding affinity and 16 drug-like indexes.

### Comparing ligands in RNALID with FDA-approved drugs and ligands without bioactivities

We collected 7,753 FDA-approved drugs from DrugBank database^30-34^, and 42,234 ligands without bioactivities from PubChem database (https://pubchem.ncbi.nlm.nih.gov/). The ligands without bioactivities were named as control ligands. For these ligands, we calculated their chemical properties, 2D and 3D structures and drug-like indexes.

To compare the chemical properties of ligands in RNALID with the FDA drugs and the control ligands, we performed PCA analysis on the cheminformatic parameters. As shown in Figure 5A, many MV ligands and SM ligands in RNALID are separated from FDA drugs and control ligands. The PC1 differences were evaluated by two-tailed MW-U tests (Figure 5A). When the RNALID ligands were grouped into solely binding with RNA and binding with RNA&protein, we found that ligands solely binding with RNA are significantly different from the FDA drugs and the control ligands in PC1. When the ligands from RNALID were divided into types of binding with vRNA and mRNA, they both were significantly different from the FDA drugs and the control ligands in PC1. When the ligands were grouped into types of rep RNA and non-rep RNA, they both were significantly distinctive from the FDA drugs and the control ligands in PC1.

**Figure 5.**
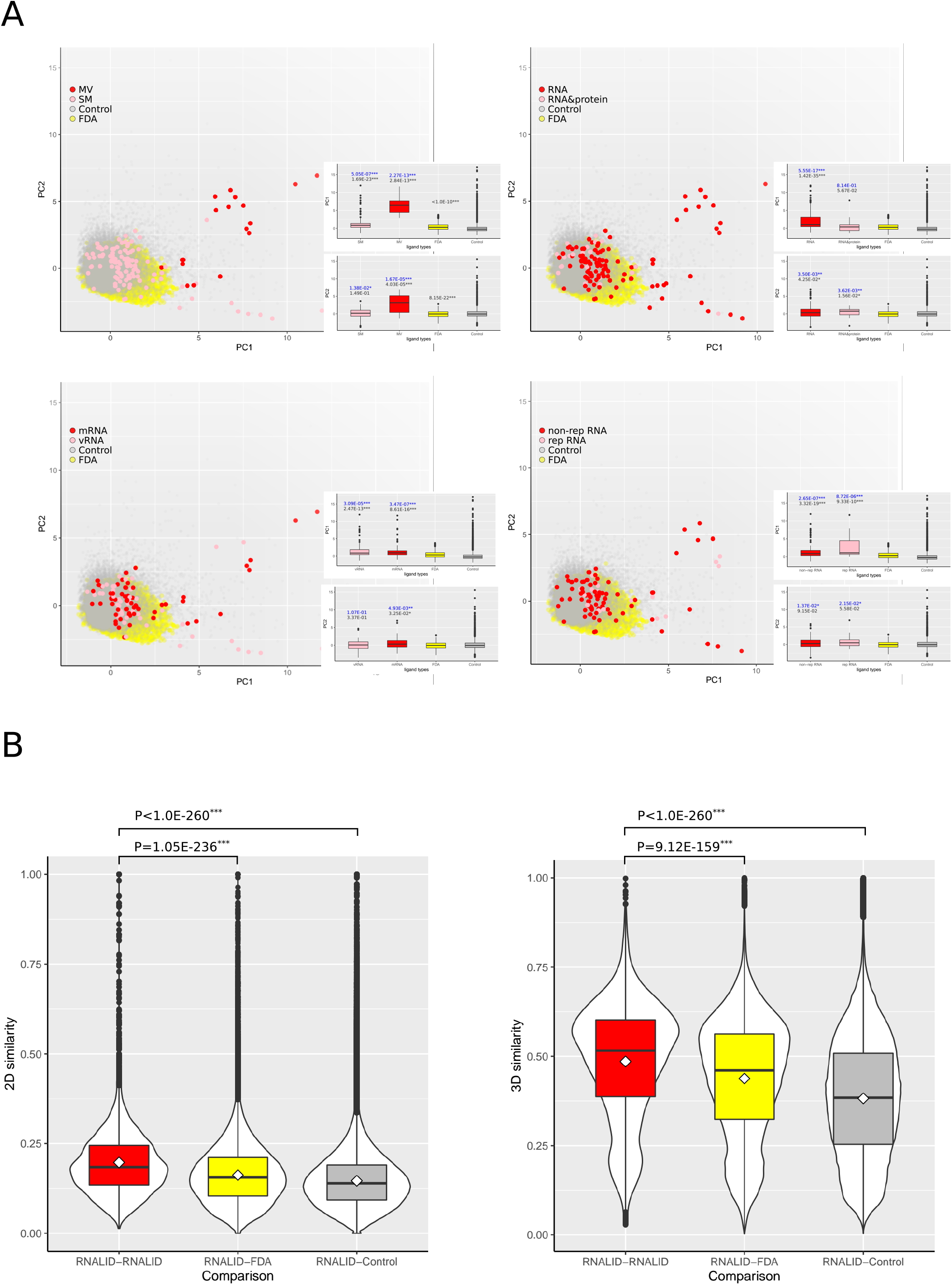
Comparing chemical characters of ligands in RNALID, FDA approved drugs and ligands without bioactivities. **(A)** Principal component analysis (PCA) results of chemical characters of ligands from three sources. Two-tailed MW-U tests were performed to evaluate the difference of them on PC1. **(B)** Comparing ligand structure similarity. Single-tail Student’s t-test was used to evaluate the differences of ligands from three sources.

Further analysis is to compare cheminformatic parameters for ligands in RNALID, the FDA drugs, and the control ligands by two-tailed MW-U tests. As shown in Table S6, only two cheminformatic parameters, Fsp3 and nStereo are not significantly different between the RNALID ligands and the control ligands. Only one cheminformatic parameter, LogP, is not significantly different between the RNALID ligands and the FDA drugs.

We further compared the 2D and 3D structures of RNALID ligands, FDA drugs, and control ligands. The intra-similarity of RNALID ligands was compared with inter-similarity with FDA drugs and control ligands. As shown in Figure 5B, the intra-similarities of RNALID ligands are significantly higher than the inter-similarity with the FDA drugs or the control ligands, indicating that the RNALID ligands are different from both the FDA drugs and the control ligands in structures.

The drug-likeness of the RNALID ligands was compared with the FDA drugs and the control ligands. We found that QED scores of *CYP2D6_I, CYP3A4_I, HERG*, and *LIVER* are not significantly different between the RNALID ligands and the control ligands (Table S7). Further comparison on the RNALID ligands and the FDA drugs indicated that *CYP1A2_I, CYP2D6_I*, and *LIVER* are not significantly different between these two groups of ligands (Table S7).

## DISCUSSION

RNAs have been considered as potential drug targets ^3, 36^. However, the properties of RNA-ligand interactions are still poorly understood which has hindered the design of RNA-targeting drugs. Recently, several databases collecting RNA-ligand interactions have been established, whereas most of them have limited information to comprehensively reflect the characteristics of RNA-ligand interactions. To date, there are no studies carrying comprehensive analysis on the characteristics of RNA-ligand interactions by dividing ligands into different types under varieties of classification principles, taken both ligand and target characters into consideration.

This study established a database, RNALID, containing the largest number of experimentally validated RNA-ligand interactions and the most comprehensive information attached with them. In RNALID, 41.48% ligands have novel structures comparing to the current databases. The novel ligands in RNALID will provide important information for characterizing RNA-ligand interactions (Table S8). When the ligands in RNALID are divided into many types using variety of classification principles, the ligands in different groups show various characteristics. As shown in Figure 6, MV ligands exhibited significantly higher values of 9 cheminformatic parameters comparing to SM ligands, indicating that SM ligands are more approaching the drug-like rules as illustrated in the study of protein-binding ligands^5^. Meanwhile, mRNA ligands exhibited more significantly higher values of drug-like indexes (11/28) comparing to vRNA ligands, which indicates that ligands targeting on mRNA may be a good candidate for drug development. Ligands binding to short-repeat RNAs (rep RNA) were the only type of ligands that exhibited significantly higher structural conservative comparing with its opposite ligand type, non-rep RNA ligands both in 2D and 3D structure comparisons. To discover if ligands belonging to one group preferred to have characteristics of other groups, we additionally analysed the enrichment of ligands in multiple groups. As shown in Figure S11, MV ligands are enriched with ligands binding with rep RNA and ligands binding with tRNA while SV ligands are enriched with ligands binding with vRNA.

**Figure 6.**
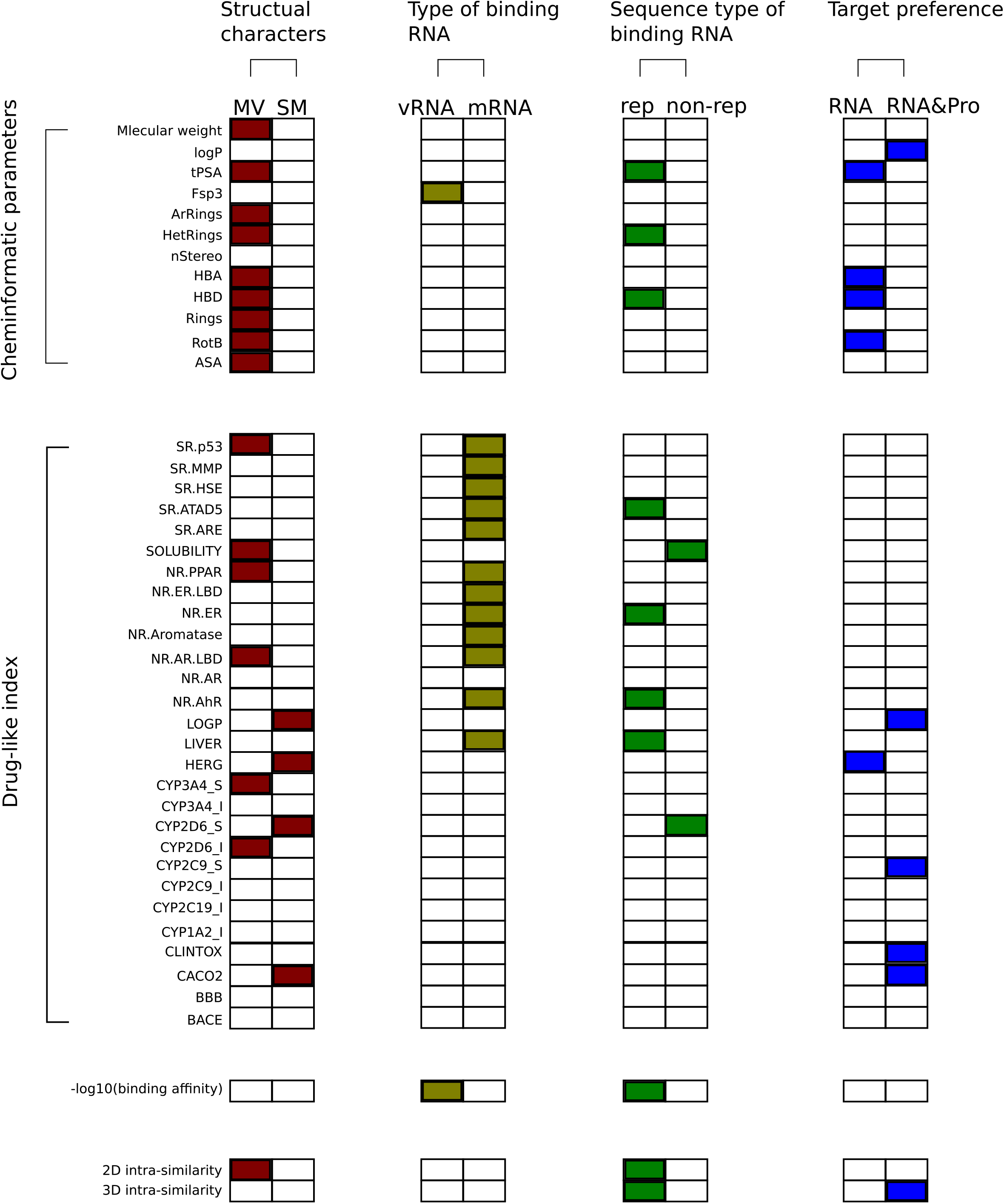
Comparing four pairs of ligands in RNALID on cheminformatic parameters, drug-like index, binding affinity, and structure similarity. **“**MV” represents multivalent ligands and “SM” represents small molecular. “vRNA” represents ligands binding with mRNA from virus and “mRNA” represents ligands binding with mRNA from cellular. “rep” represents ligands binding with RNA with repeat sequences and “non-rep” represents ligands binding with RNA without repeat sequence. “RNA” and “RNA & pro” represent ligands only binding with RNA and ligands binding with both RNA and protein, respectively. The squares in red, yellow, green and blue represent types of ligands with significantly (P-value < 0.05) higher values than others.

Drug-likeness is a key factor in drug development^37^. It has been proved that more than 40% of compounds failed to be developed into drugs due to drug-like properties^38^. Thus, it is important to discover the relationship among drug-like indexes, RNA-ligand binding affinity and ligand properties. In this study, 2D and 3D structural characteristics, chemical properties of ligands, and the relationship between ligand binding affinity and 28 drug-like indexes were analyzed for ligands grouped under various classification principles. We found that the drug-like indexes *CACO2, NR*.*AhR, SOLUBILITY* and *SR*.*HSE* of ligands were the most frequently related with binding affinity of ligands from different types. In most of the cases, CACO2 cell model was used to validate the capability of drugs to go through human Intestinal epithelium^39^. Besides, previous studies have indicated the close relationship between *CACO2* cell activities and protein targets^40-46^, for example, antioxidant agent can lead to the morphological changes and cell viability for Vero and Caco-2 cell line^40^, exhibiting close relationship between caco-2 cell activity and oxidation circumstances^41, 42^.

Studies have revealed that many virus genomes contain structures or sequence motifs for binding ligands^47^. However, no studies have been compared the ligands binding with virus mRNA (vRNA) to ligands binding with mRNAs from cellular organisms^5, 17, 18, 48, 49^. The analysis in this study indicated that ligands binding to mRNAs and vRNAs exhibited significant difference in binding affinities, chemical characters and co-relationships with drug-like indexes (Figure S11). Additionally, we found that vRNA-binding ligands were enriched with ligands binding with non-rep RNA. In comparison, ligands binding with mRNAs from cellular organisms were enriched with ligands binding to rep RNAs (Figure S11). A potential explanation on the differences is the difference of ligand structures. vRNAs prefer to bind with SM ligands while mRNAs from cellular organisms prefer to binding with MV ligands. More efforts are required to reveal the biological mechanisms underlying the ligand binding preferences of vRNA and mRNAs in cellular organisms.

To distinguish the properties of ligands in RNALID with ligands not binding with RNA, we compared RNALID with FDA approved protein-binding drugs and control ligands without bioactivity in several respects. Two drug-like indexes, *LIVER* and *CYP2D6_I*, are not significantly different between RNALID ligands and FDA drugs, and between RNALID ligands and the control ligands. Further study is required to detect the underlying reasons.

In conclusion, this study established the RNALID, a comprehensive database collecting RNA-ligand interactions. By analyzing RNALID database, the signatures of different types of ligands in chemical characters, spatial properties, binding affinities, structural conservation were described. Further drug-likeness evaluation provided clues to discover ligands with high potential to be drugs.

## Supporting information

RNA_drug_suppl210914.docx

## DATA AVAILABILITY

RNALID was available at http://121.46.19.89/RNALID/html/index.html#/database

## SUPPLEMENTARY DATA

Supplementary Data are available at NAR online.

## ACKNOWLEDGEMENT

The work was partly funded by the National Key R&D Program of China (2020YFB0204803), the Natural Science Foundation of China (81801132, 81971190, 61772566), Guangdong Key Field R&D Plan (2019B020228001 and 2018B010109006), Introducing Innovative and Entrepreneurial Teams (2016ZT06D211), Guangzhou S&T Research Plan (202007030010), and Mater Foundation (YH.Y).

## FIGURES LEGENDS

### Supplementary materials

RNA_drug_suppl210914.docx

## REFERENCES

1. Cheng, A. C., et al., Structure-based maximal affinity model predicts small-molecule druggability. Nature biotechnology 2007, 25 (1), 71–75.

2. Harrow, J., et al., GENCODE: the reference human genome annotation for The ENCODE Project. Genome Res 2012, 22 (9), 1760–74.

3. Barry, G., Integrating the roles of long and small non-coding RNA in brain function and disease. Mol Psychiatry 2014, 19 (4), 410–6.

4. Nord, A. S.; West, A. E., Neurobiological functions of transcriptional enhancers. Nat Neurosci 2020, 23 (1), 5–14.

5. Warner, K. D., et al., Principles for targeting RNA with drug-like small molecules. Nat Rev Drug Discov 2018, 17 (8), 547–558.

6. Desterro, J., et al., Targeting mRNA processing as an anticancer strategy. Nat Rev Drug Discov 2020, 19 (2), 112–129.

7. Han, T., et al., Anticancer sulfonamides target splicing by inducing RBM39 degradation via recruitment to DCAF15. Science 2017, 356 (6336).

8. Webb, T. R., et al., The development and application of small molecule modulators of SF3b as therapeutic agents for cancer. Drug Discov Today 2013, 18 (1-2), 43–9.

9. Jin, K. T., et al., Roles of lncRNAs in cancer: Focusing on angiogenesis. Life Sci 2020, 252, 117647.

10. Mitash, N., et al., The Role of MicroRNA in the Airway Surface Liquid Homeostasis. Int J Mol Sci 2020, 21 (11).

11. Song, W., et al., The Emerging Roles of Long Noncoding RNAs in Bone Homeostasis and Their Potential Application in Bone-Related Diseases. DNA Cell Biol 2020, 39 (6), 926–937.

12. Toni, L., et al., Dysregulated micro-RNAs and long noncoding RNAs in cardiac development and pediatric heart failure. Am J Physiol Heart Circ Physiol 2020, 318 (5), H1308–H1315.

13. Petrescu, G. E. D., et al., MicroRNA based theranostics for brain cancer: basic principles. J Exp Clin Cancer Res 2019, 38 (1), 231.

14. Costales, M. G., et al., A Designed Small Molecule Inhibitor of a Non-Coding RNA Sensitizes HER2 Negative Cancers to Herceptin. J Am Chem Soc 2019, 141 (7), 2960–2974.

15. Serganov, A., et al., Coenzyme recognition and gene regulation by a flavin mononucleotide riboswitch. Nature 2009, 458 (7235), 233–7.

16. Kumar Mishra, S.; Kumar, A., NALDB: nucleic acid ligand database for small molecules targeting nucleic acid. Database (Oxford) 2016, 2016.

17. Morgan, B. S., et al., Discovery of Key Physicochemical, Structural, and Spatial Properties of RNA-Targeted Bioactive Ligands. Angew Chem Int Ed Engl 2017, 56 (43), 13498–13502.

18. Morgan, B. S., et al., Insights into the development of chemical probes for RNA. Nucleic Acids Res 2018, 46 (16), 8025–8037.

19. Morgan, B. S., et al., R-BIND: An Interactive Database for Exploring and Developing RNA-Targeted Chemical Probes. ACS Chem Biol 2019, 14 (12), 2691–2700.

20. Velagapudi, S. P., et al., Sequence-based design of bioactive small molecules that target precursor microRNAs. Nat Chem Biol 2014, 10 (4), 291–7.

21. O’Boyle, N. M., et al., Open Babel: An open chemical toolbox. J Cheminform 2011, 3, 33.

22. Gilson, M. K., et al., BindingDB in 2015: A public database for medicinal chemistry, computational chemistry and systems pharmacology. Nucleic Acids Res 2016, 44 (D1), D1045–53.

23. Liu, T., et al., BindingDB: a web-accessible database of experimentally determined protein-ligand binding affinities. Nucleic Acids Res 2007, 35 (Database issue), D198–201.

24. Yang, Y., et al., SPOT-Ligand: Fast and effective structure-based virtual screening by binding homology search according to ligand and receptor similarity. J Comput Chem 2016, 37 (18), 1734–9.

25. Hu, J., et al., LS-align: an atom-level, flexible ligand structural alignment algorithm for high-throughput virtual screening. Bioinformatics 2018, 34 (13), 2209–2218.

26. Fan, C., et al., DStruBTarget: Integrating Binding Affinity with Structure Similarity for Ligand-Binding Protein Prediction. J Chem Inf Model 2020, 60 (1), 400–409.

27. Ying Song, S. Z., Zhangming Niu, Zhang-Hua Fu, Yutong Lu, Yuedong Yang, Communicative Representation Learning on Attributed Molecular Graphs. Proceedings of the Twenty-Ninth International Joint Conference on Artificial Intelligence (IJCAI-20 2019, (IJCAI-20), 7.

28. Delaney, J. S., ESOL: estimating aqueous solubility directly from molecular structure. J Chem Inf Comput Sci 2004, 44 (3), 1000–5.

29. Mobley, D. L.; Guthrie, J. P., FreeSolv: a database of experimental and calculated hydration free energies, with input files. J Comput Aided Mol Des 2014, 28 (7), 711–20.

30. Knox, C., et al., DrugBank 3.0: a comprehensive resource for ‘omics’ research on drugs. Nucleic Acids Res 2011, 39 (Database issue), D1035–41.

31. Law, V., et al., DrugBank 4.0: shedding new light on drug metabolism. Nucleic Acids Res 2014, 42 (Database issue), D1091–7.

32. Wishart, D. S., et al., DrugBank 5.0: a major update to the DrugBank database for 2018. Nucleic Acids Res 2018, 46 (D1), D1074–D1082.

33. Wishart, D. S., et al., DrugBank: a knowledgebase for drugs, drug actions and drug targets. Nucleic Acids Res 2008, 36 (Database issue), D901–6.

34. Wishart, D. S., et al., DrugBank: a comprehensive resource for in silico drug discovery and exploration. Nucleic Acids Res 2006, 34 (Database issue), D668–72.

35. Lipinski, C. A., Lead-and drug-like compounds: the rule-of-five revolution. Drug Discov Today Technol 2004, 1 (4), 337–41.

36. Blount, K. F., et al., Novel riboswitch-binding flavin analog that protects mice against Clostridium difficile infection without inhibiting cecal flora. Antimicrob Agents Chemother 2015, 59 (9), 5736–46.

37. Agoni, C., et al., Druggability and drug-likeness concepts in drug design: are biomodelling and predictive tools having their say? J Mol Model 2020, 26 (6), 120.

38. Hoffman, A. S., Hydrogels for biomedical applications. Adv Drug Deliv Rev 2002, 54 (1), 3–12.

39. Gil-Ramirez, A., et al., Modulation of cholesterol-related gene expression by ergosterol and ergosterol-enriched extracts obtained from Agaricus bisporus. Eur J Nutr 2016, 55 (3), 1041–57.

40. Hassan, S. E., et al., Endophytic actinomycetes Streptomyces spp mediated biosynthesis of copper oxide nanoparticles as a promising tool for biotechnological applications. J Biol Inorg Chem 2019, 24 (3), 377–393.

41. Hou, Y., et al., Transcriptomic responses of Caco-2 cells to Lactobacillus rhamnosus GG and Lactobacillus plantarum J26 against oxidative stress. J Dairy Sci 2019, 102 (9), 7684–7696.

42. Katayama, S., et al., Antioxidative stress activity of oligophosphopeptides derived from hen egg yolk phosvitin in Caco-2 cells. J Agric Food Chem 2006, 54 (3), 773–8.

43. Lindemann, G., et al., Limited heat-shock protein 72 induction in Caco-2 cells by L-glutamine. Digestion 2001, 64 (2), 81–6.

44. Yu, M., et al., Aryl Hydrocarbon Receptor Activation Modulates Intestinal Epithelial Barrier Function by Maintaining Tight Junction Integrity. Int J Biol Sci 2018, 14 (1), 69–77.

45. Gu, S., et al., Activation of membrane androgen receptors in colon cancer inhibits the prosurvival signals Akt/bad in vitro and in vivo and blocks migration via vinculin/actin signaling. Mol Med 2011, 17 (1-2), 48–58.

46. Swank, G. M., et al., Effect of acute-phase and heat-shock stress on apoptosis in intestinal epithelial cells (Caco-2). Crit Care Med 1998, 26 (7), 1213–7.

47. Hermann, T., Small molecules targeting viral RNA. Wiley Interdiscip Rev RNA 2016, 7 (6), 726–743.

48. Rzuczek, S. G., et al., Studying a Drug-like, RNA-Focused Small Molecule Library Identifies Compounds That Inhibit RNA Toxicity in Myotonic Dystrophy. ACS Chem Biol 2015, 10 (12), 2706–15.

49. Aboul-ela, F., Strategies for the design of RNA-binding small molecules. Future Med Chem 2010, 2 (1), 93–119.

