## Supplementary material for "Characterizing RNA-binding Ligands on Structures, Chemical Information, Binding Affinity and Drug-likeness": RNA_drug_suppl210914.docx

**Supplementary Note**

**Method**

**Chemical similarities between ligands**

Chemical similarity is calculated as the Euclidean distance between two ligands in the space of cheminformatic parameters as done before^1^. First, for each of the 12 cheminformatic parameters, the mean value (E) and standard deviation (STD) is calculated for all SM ligands in RNALID database. Then, for the query ligand, each of the 12 cheminformatic parameters was calculated and normalized using E and STD values of that chemical characteristic, as shown in Equation 1.

$X_{norm}=\frac{(X-E)}{STD}$ (Equation 1),

where *X* is the value of cheminformatic parameters, *X_norm_* is the normalized value, *E* and *STD* are the mean and standard deviation among all SM ligands in RNALID database, respectively.

The chemical similarity between two ligands is evaluated by the Euclidean distance calculated as Equation 2.

$ED=\sqrt{\sum_{i=1}^{12} {(X_{i1}-X_{i2})}^{2}}$ (Equation 2),

where *ED* is the Euclidean distance between two ligands in the space of cheminformatic parameters, and *X_i1_* and *X_i2_* is the normalized value of the *i*th cheminformatic parameters (*X_i_*) of the ligand *1* and the ligand *2*, respectively.

**Clustering ligands by RDKit**

The ligand structure clustering was carried out by RDKit (http://www.rdkit.org/) based on fingerprint. First, the MorganFingerprint was generated for each ligand by the function of *GetMorganFingerprintAsBitVect()* in the module of AllChem. For each pair of ligand fingerprints, the similarity was calculated by the function of *BulkTanimotoSimilarity()*. Then, function of *Butina.ClusterData()* was used to cluster the ligands according to their fingerprint similarities. Finally, the first ligand of each cluster was picked up as structure representative of this cluster, and these structure representatives were further used for enrichment analysis among ligand types.

**Ligand spatial property**

The calculation of principal moment of interias (PMI) were performed by RDKit software. *AddHs()* function of RDKit was used to add hydrogen atoms for each ligand. Then, 20 energy conformations were generated by the function of *EmbedMultipleConfs()* in AllChem module, which were further optimized by the function of *MMFFOptimizeMoleculeConfs()*. For each optimized conformation, the PMI indexes (*I_1_*, *I_2_* and *I_3_*) were calculated by the function of *Descriptors3D.PMI1()*, *Descriptors3D.PMI2()* and *Descriptors3D.PMI3()*, separately. The normalized PMI indexes, *npr1* and *npr2* were calculated as *I_1_*/*I_3_* and *I_2_*/*I_3_*, respectively. Finally, for each ligand, its *npr1* and *npr2* were calculated by *Equation* 3.

${npr}_{1,2}=\frac{\sum\mathrm{npr}_{i}*e^{\frac{-E_{i}}{K_{B}*T}}}{\sum e^{\frac{-E_{i}}{K_{B}*T}}}$ (Equation 3),

where *npr*_i_ represent value of *i* th low-energy conformation of the ligand, *K_B_* is the Boltzmann constant (0.001986 kcal/(mol*K)), *T* is the temperature (298 *K*), and $E_{i}$ is the energy of the *i* th conformation of the ligand. When plotting *npr1* and *npr2* of each ligand in the triangular graph, the three endpoints of the triangle represent the shapes of rod- (0,1), sphere- (1,1) or disc-like (0.5, 0.5)^2^.

To quantitively access the spatial property of one type of ligands (using rod as an example as below), the distances between rod endpoint (0, 1) to ligands were calculated. Then, the set of distances of this ligand type was compared with another ligand type. The statistical significances were evaluated by a two-tail MW-U test. P-value<0.05 indicates that ligands in one type is significantly more approaching to one shape than to others. Accumulative distributions of distance between ligand points and endpoints were plotted to detect the spatial difference by visual.

**RNA similarity**

RNA similarity includes RNA secondary structure and RNA sequence similarities. The RNA secondary structure are predicted by the module of RNAfold in the ViennaRNA Package^3^ by using the RNA sequence information. The secondary structure similarity was calculated by SimTree (http://bioinfo.cs.technion.ac.il/SimTree/Abstract.html), and the similarity values are ranging from 0 to 1, with a higher value denotes a higher structure similarity. The sequence similarity of RNA was calculated by BLAST^4^. E-value obtained from BLAST was used to evaluate the sequence similarity between RNAs. Lower E-value means higher sequence similarity.

**Result**

**User interface of RNALID**

The interface of RNALID allows users to browsing, searching and downloading data. As shown in Figure S1A, RNALID includes three modules, “Ligand search”, “RNA search” and “RNA-ligand interaction”. “Ligand search” is to search RNAs potentially binding with the query ligand. By using ligands in RNALID as templates (reference ligands), RNAs binding with the reference ligands that are structurally or chemically similar to the query ligands will be shown as results. Users are allowed to submit query ligands in several manners including: (1) uploading a file including structures (in the format of SMILES, 2D-SDFs or 3D-SDFs) of multiple ligands (Figure S1B and C); (2) drawing a molecule in a java plug-in (Figure S1B); and (3) using a DrugBank ID^30-34^ (in the format of “DBXXXX”). After clicking on “Submit”, RNALID will use the structure of the query ligand to predict RNAs binding with it (Figure S1C). The top three results (ranked according to similarity between reference ligand and the query) of the query will be provided in the webpage. All results can be downloaded in a zipped package (Figure S1D). For each RNA that was predicted as binding with the query ligand, RNALID provided name of the RNA, the 2D and 3D structure similarity scores between the query and the reference ligands, chemical similarity score between the query and the reference ligands, binding affinity between the predicted RNA and the reference ligands, and the associated diseases of the reference ligands.

“RNA search” module is to find ligands potentially binding with the query RNA, as well as RNAs having similar sequence or secondary structure with the query RNA. Users are allowed to input RNA information in the form of sequence or secondary structure (parentheses representation) as shown in Figure S1E. Meanwhile, users are allowed to do batch searching by uploading a file containing information on multiple RNAs. If the input is RNA sequence, RNALID will use BLAST software^36^ to align the sequences of query and reference RNAs. If the sequence alignment E-score is less than 0.05, RNALID will provide the ligands binding with the reference RNAs(Figure S1F). When only RNA secondary structure is input, RNALID will give results solely based on RNA secondary structure similarity calculated by SimTree module in the ViennaRNA package^37^. Furthermore, the user can simultaneously input information of a ligand as well as an RNA sequence or secondary structure to search if they are potentially interacted with each other. RNALID will provide the top three most similar ligands (Figure S1D), similar RNAs (Figure S1F), and the most possible interactions as shown in Figure S1G. If the input exhibited 100% similarity with the ligands or RNA in RNALID, the output interactions will be marked as red.

**Comparing RNALID to other databases on chemical and structure properties**

The twelve cheminformatic parameters of ligands were calculated by RDKit as shown in Table S2. The values of these cheminformatic parameters of ligands were used to cluster the ligands by Principle component analysis (PCA). As shown in Figure S3, two groups of ligands are not clustered together, which represent novel chemical properties of newly collected ligands in RNALID comparing to R-BIND database. To further study the chemical properties of ligands in RNALID, we calculated the chemical similarity scores among ligands. The newly collected ligands in RNALID represent lower similarity comparing to the ligands in R-BIND. 94.64% of RNALID ligands are found with pair-wised inter- and intra- chemical similarity scores lower than 0.25 simultaneously (lower quartile). In comparison, only 57% ligands in R-BIND have pair-wised chemical similarity score lower than 0.25.

The structure properties of ligand include two-dimensional (2D) structure and three-dimensional (3D) structure. The pair-wised 2D or 3D structure similarities of ligands were calculated by tanimoto coefficient as used by SpotLigand^10^ and LSalign^12^, respectively. Ligands in RNALID database exhibited higher 2D structure novelty than R-BIND database (Figure S4A), but not in 3D structure (Figure S4B). As shown in Figure S4, “*X*” axis is the 2D structure similarity calculated for ligands within the same database (intra-database similarity), and “*Y”* axis is the 2D structure similarity calculated for ligands from different databases (inter–database similarity). For each ligand, the top 2D structure similarity scores respectively from inter comparison and intra comparison were used to evaluate structure diversity. As shown in Figure S4, 21.4% newly collected ligands in RNALID are found with 2D structure similarity scores lower than 0.4 in both intra and inter structure comparison. Only 12.8% ligands from R-BIND have 2D structure similarity scores lower than 0.4 in both intra and inter structure comparison.

**PMI analysis of FDA drugs, the control ligands and RNALID ligands**

Spatial properties of FDA drugs, the control ligands and RNALID ligands were evaluated as shown in Figure S10. It can be seen that FDA drugs and control ligands were always significantly different from each other in distance distribution from endpoints representing rod, sphere and disc. FDA drugs are more likely to be Rod than the control ligands, and they are less likely to be Sphere and Disc than the control ligands with student’s t-test P-values of 7.00E-19, 8.94E-11, and 2.02E-14, respectively. The RNALID ligands are indicated less likely to be Sphere than the control ligands with student’s t-test P-value of 9.10E-3. No significant differences were observed between FDA and RNALID ligands.

**Cross-enrichment analysis of ligand types**

Cross-enrichment analysis was employed to detect if one type of ligands is enriched with ligands belonging to another type. The enrichment was evaluated by single-tailed Fisher’s exact test. The results were shown in Figure S11. We found that ligands targeting on vRNA were significantly (P-value =1.01E-2) enriched by ligands binding with “non-rep RNA” while the ligands binding with non-virus mRNA (mRNA) were significantly enriched by ligands binding with rep-RNA (P-value of 1.31E-4). As to other RNA types, ligands binding with tRNA significantly enriched with ligands from both MV and rep-RNA groups. SM ligands were significantly enriched with vRNA ligands, while MV ligands were significantly enriched with ligands binding with tRNA and rep RNA. Notably, ligands binding solely with RNA or RNA&protein did not show enrichment in other types of ligands. Moreover, we found that ligands binding with both RNA and protein were all SM ligands.

**Strong interactions and medium interaction in RNALID database**

In RNALID, the RNA-ligand interactions were defined as strong interactions if the binding affinities were lower than 1 *μM* (1E-6 *mol/L*) and higher than 0 *nM*, and they were defined as medium interactions if the binding affinities were ranging from 1*μM* to 1 *mM* (1E-3 *mol/L*). Totally, 85 interactions were strong interactions, and 119 interactions were medium interactions. The strong interactions were involved by 47 ligands, and the medium interactions were involved by 72 ligands.

**Supplementary figures**

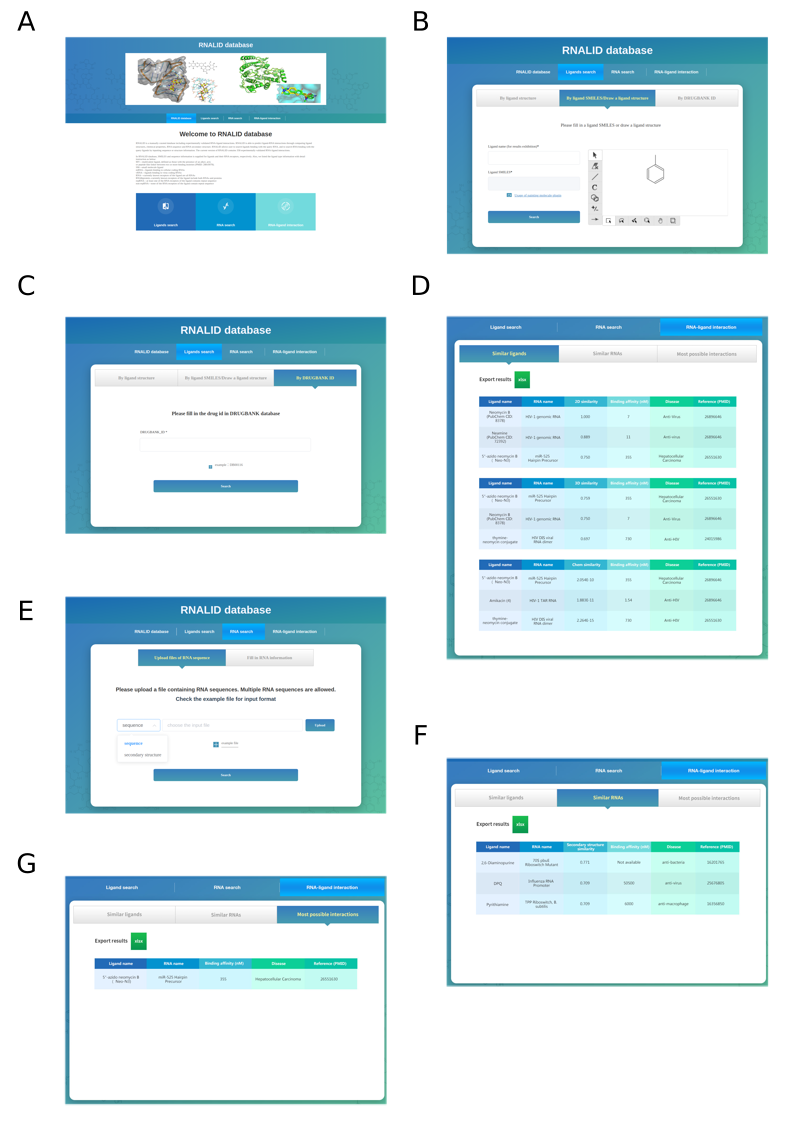

**Figure S1**. User interface of RNALID. **(A)** Home page. **(B)** The interface of RNALID for searching RNA-ligand interactions from submitted ligand. In this page, user can input ligands in the format of SMILES or drawing a ligand. **(C)** The interface of RNALID for searching RNA-ligand interaction by submitting DrugBank ID. **(D)** The output page of searching from submitted ligand or DrugBank ID. **(E)** The interface of searching RNA-ligand interactions from submitted RNA. In this page, user can input RNA in the format of sequence or secondary structure. **(F)** The output page of searching from submitted RNA. **(G)** The output page of searching from a pair of RNA and ligand.

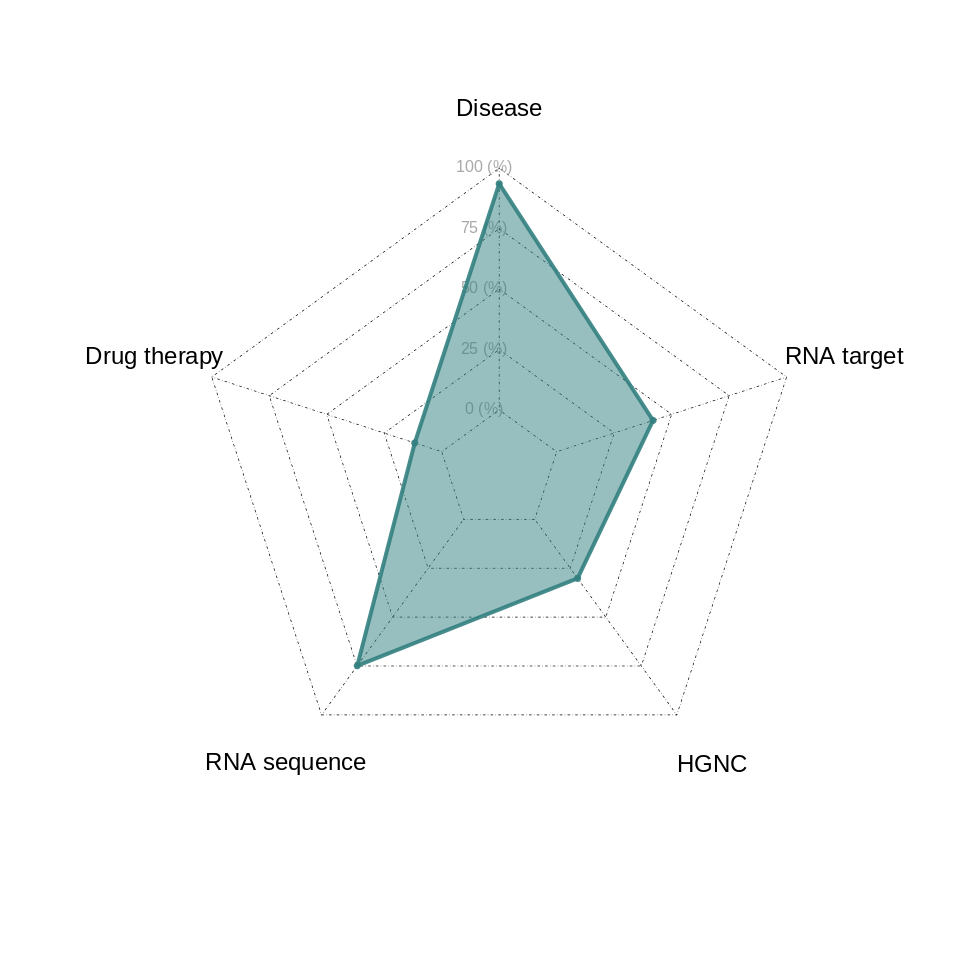

**Figure S2.** Percentages of novel information in RNALID. “Disease” means the diseases targeted by ligands in the publishing article; “Drug therapy” means clinical usage of drugs recorded in DrugBank database (https://www.drugbank.com/); “RNA sequence” means the sequence of RNA targeted by ligands in RNALID; “HGNC” means the names of human RNA; “RNA target” means direct binding or regulation non-ligand target of RNAs recorded in the references.

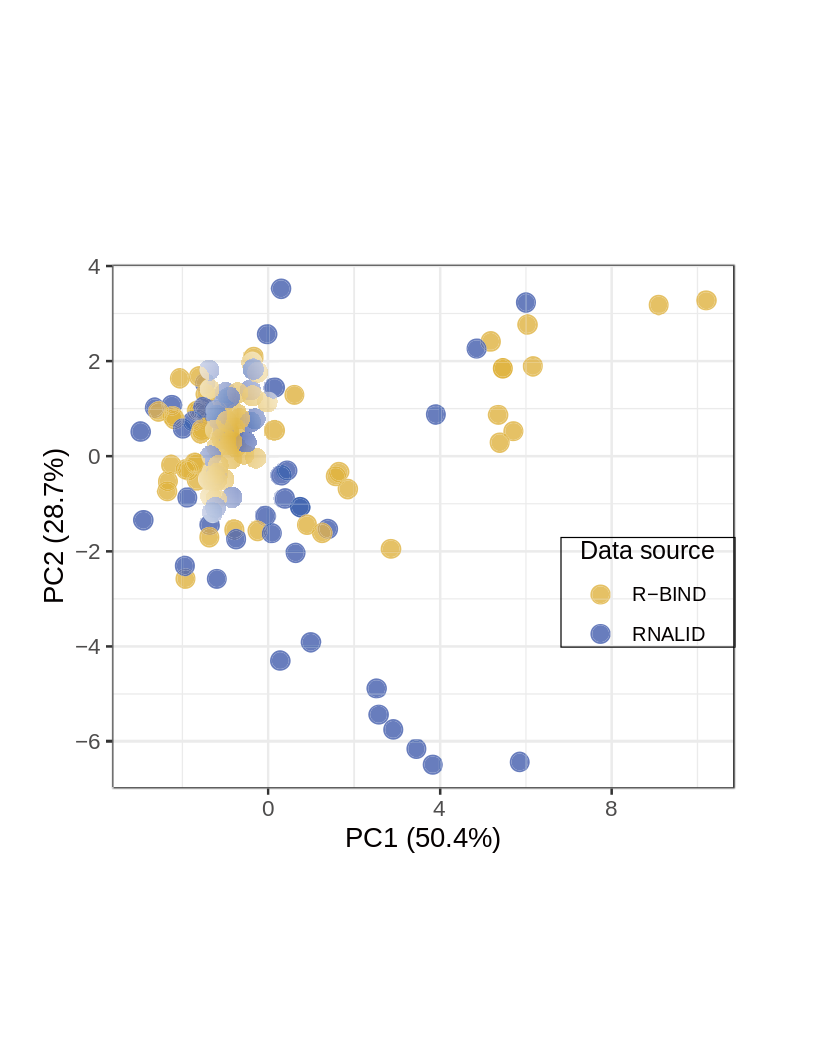

**Figure S3.** PCA analysis of 12 cheminformatic parameters of ligands in RNALID and RBIND database.

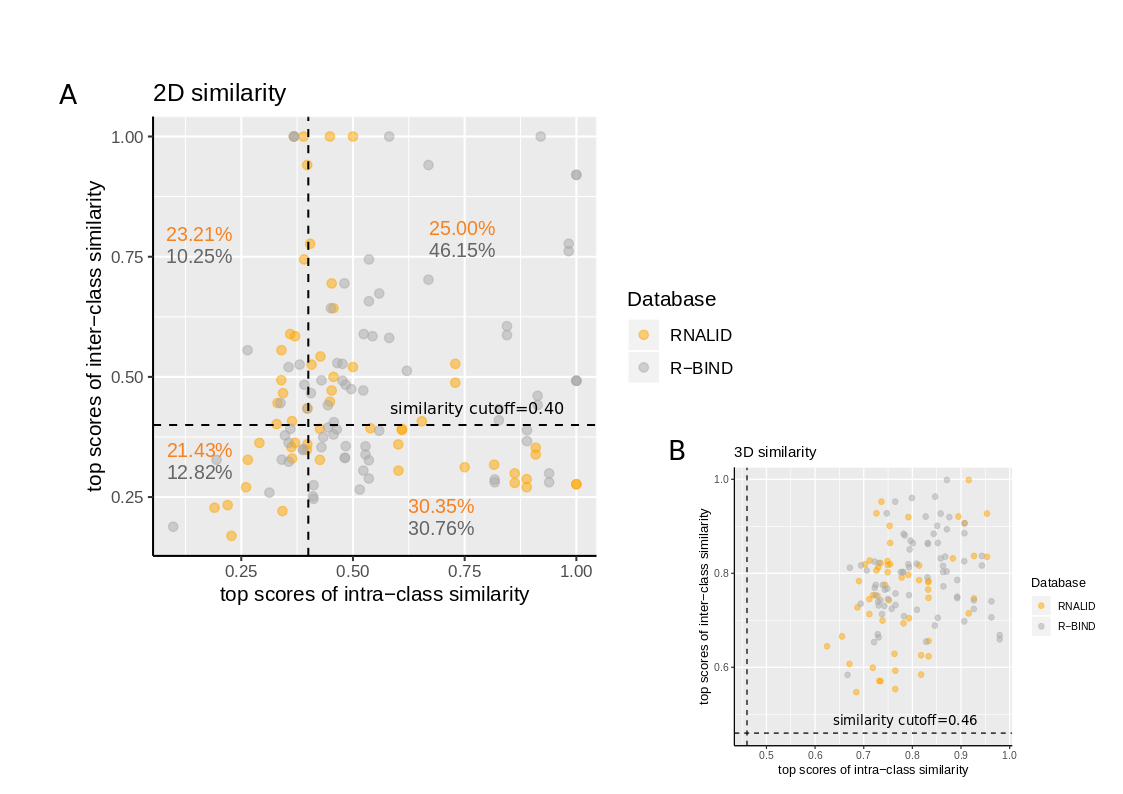
**Figure S4.** Comparing inter- and intra- structural similarity of ligands in RNALID and RBIND database. The similar cutoff was set to 0.4 for 2D similarity and 0.46 for 3D similarity, according to previous publishings^10, 12^.

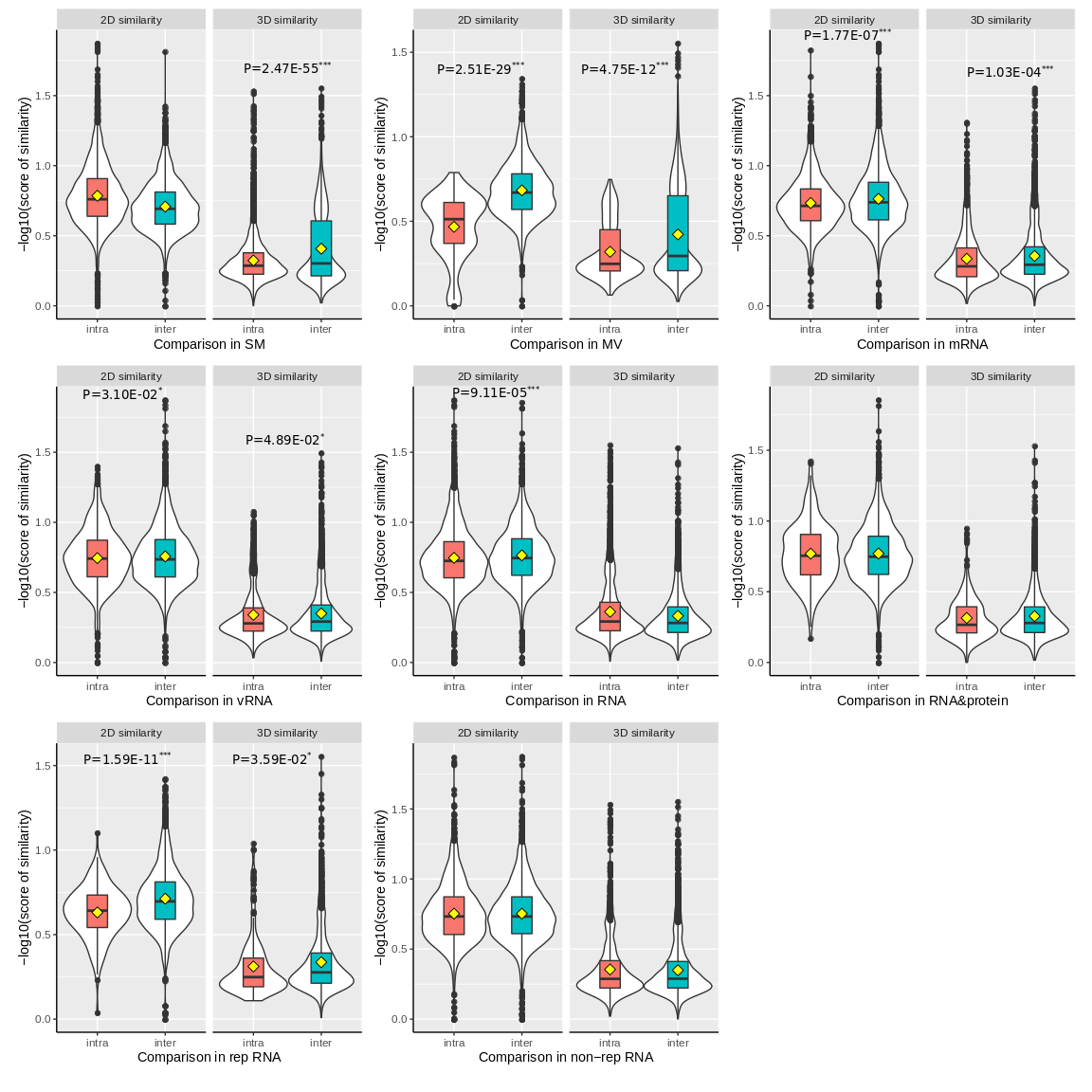

**Figure S5.** Comparison of intra- and inter- structural similarity in each type of ligands. The difference was evaluated by single-tail student’s t-test, and significant P-values indicate a higher intra-similarity than inter-similarity. “*” denotes P-value < 0.05, “**” denotes P-value < 0.01, “***” denotes P-value < 0.001.

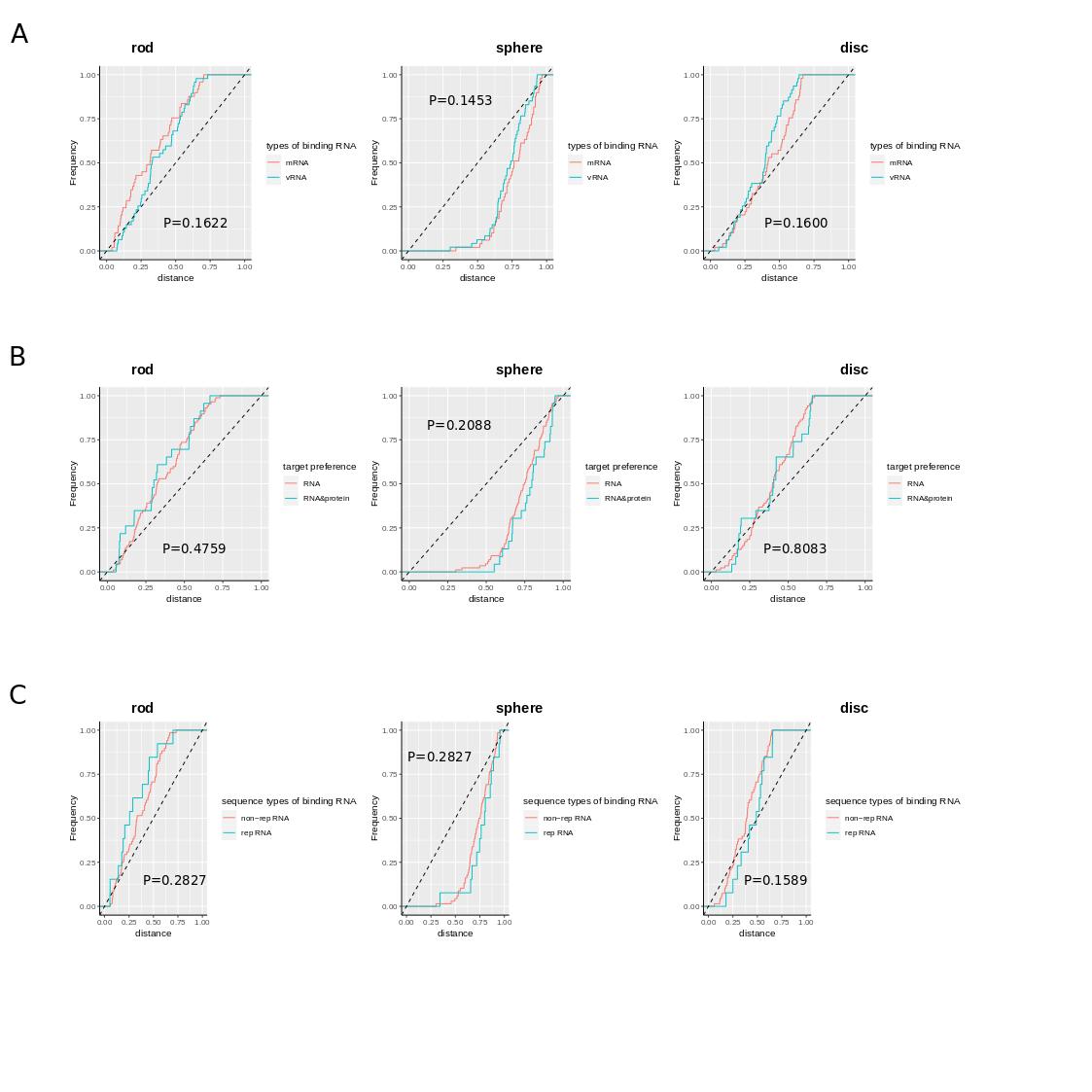

**Figure S6.** PMI accumulation distribution of different types of ligands. **(A)** ligands binding with different types of RNA; **(B)** ligands binding only with RNA and ligands binding with both RNA and protein (RNA&protein); **(C)** ligands binding with RNA with repeat sequence (rep RNA) and ligands binding with RNA without repeat sequence (non-rep RNA). Two-tailed MW-U test was used to evaluate the difference PMI accumulation distribution for different types of ligands.

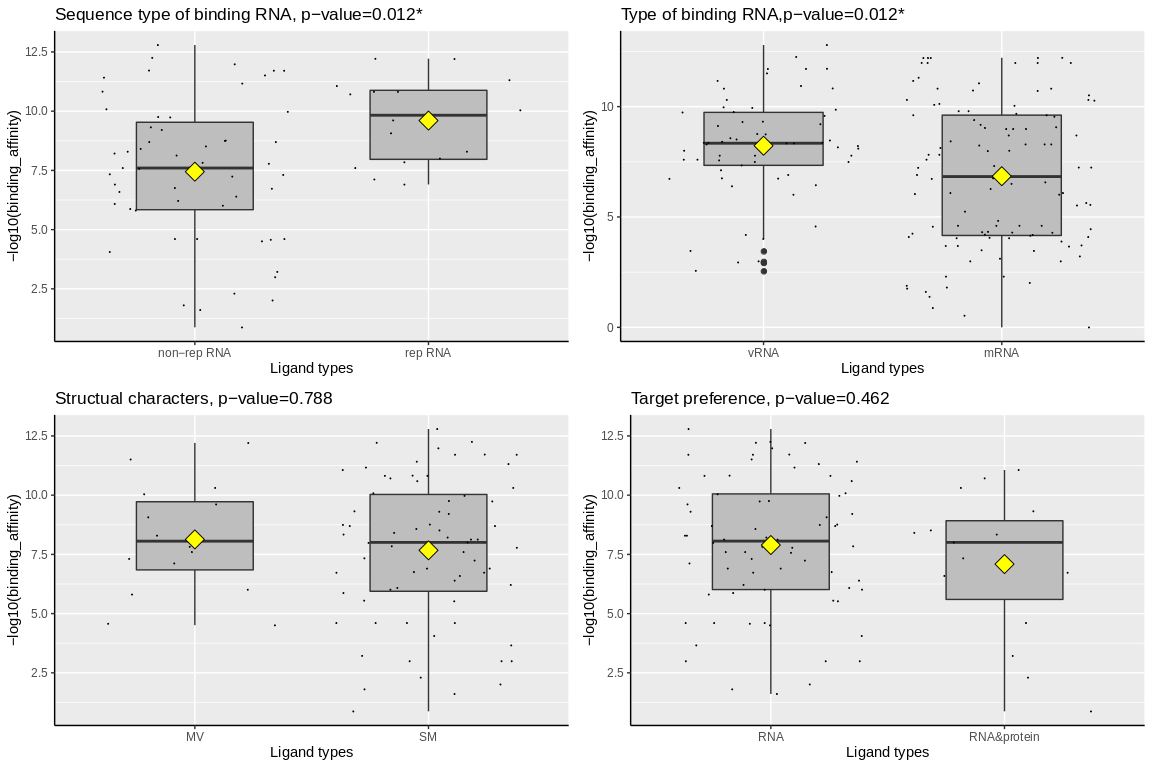

**Figure S7.** Comparison of binding affinity of different types of ligands. **(A)** Comparing binding affinity of ligands binding with RNA with repeat sequence (rep RNA) and ligands binding with RNA without repeat sequence (non-rep RNA). **(B)** Comparing binding affinity of ligands binding to cellular coding RNA (mRNA) and ligands binding virus RNA (vRNA). **(C)** Comparing binding affinity of MV ligands and SM ligands. **(D)** Comparing binding affinity of ligands only binding with RNA and ligands binding with both RNA and protein (RNA&protein). Two-tailed MW-U test was used to evaluate the difference. “*” denotes P-value < 0.05, “**” denotes P-value < 0.01, “***” denotes P-value < 0.001.

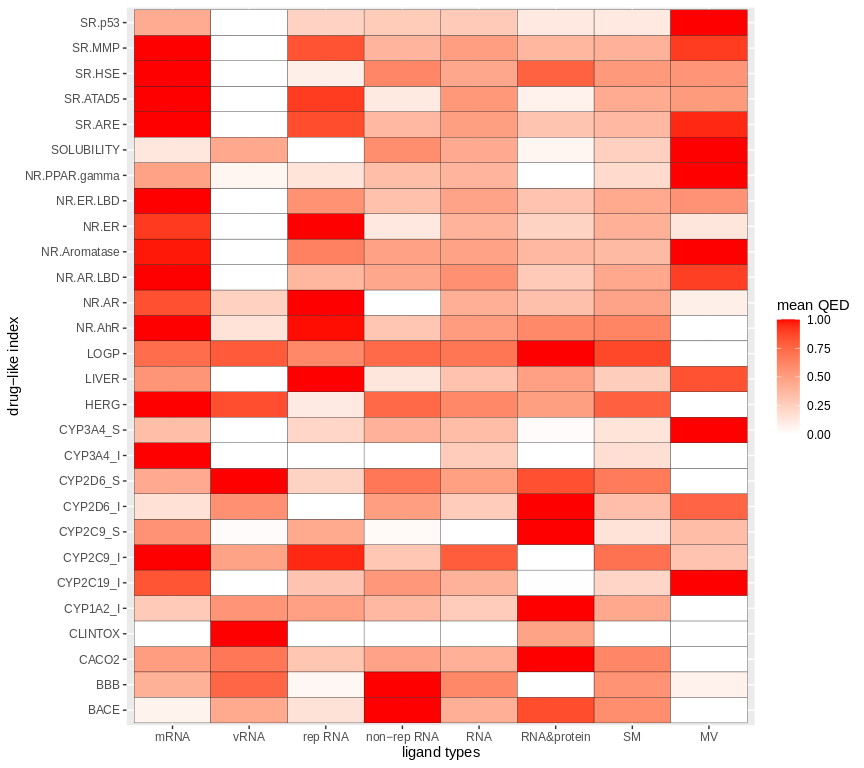

**Figure S8.** Mean of QED scores of 28 drug-like indexes and 10 ligand types.

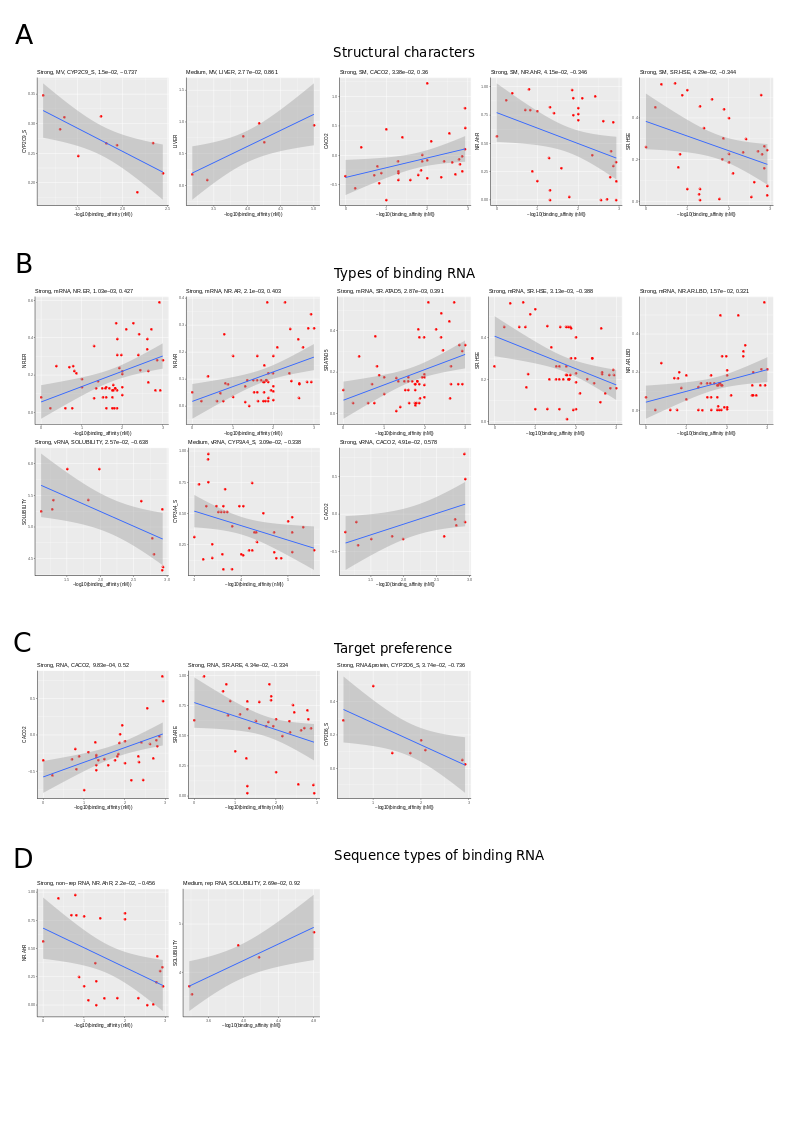

**Figure S9.** Correlation between binding affinity and QED scores of drug-like indexes. Only statistically significant results (P-value < 0.05 and PCC>0.3) are exhibited.

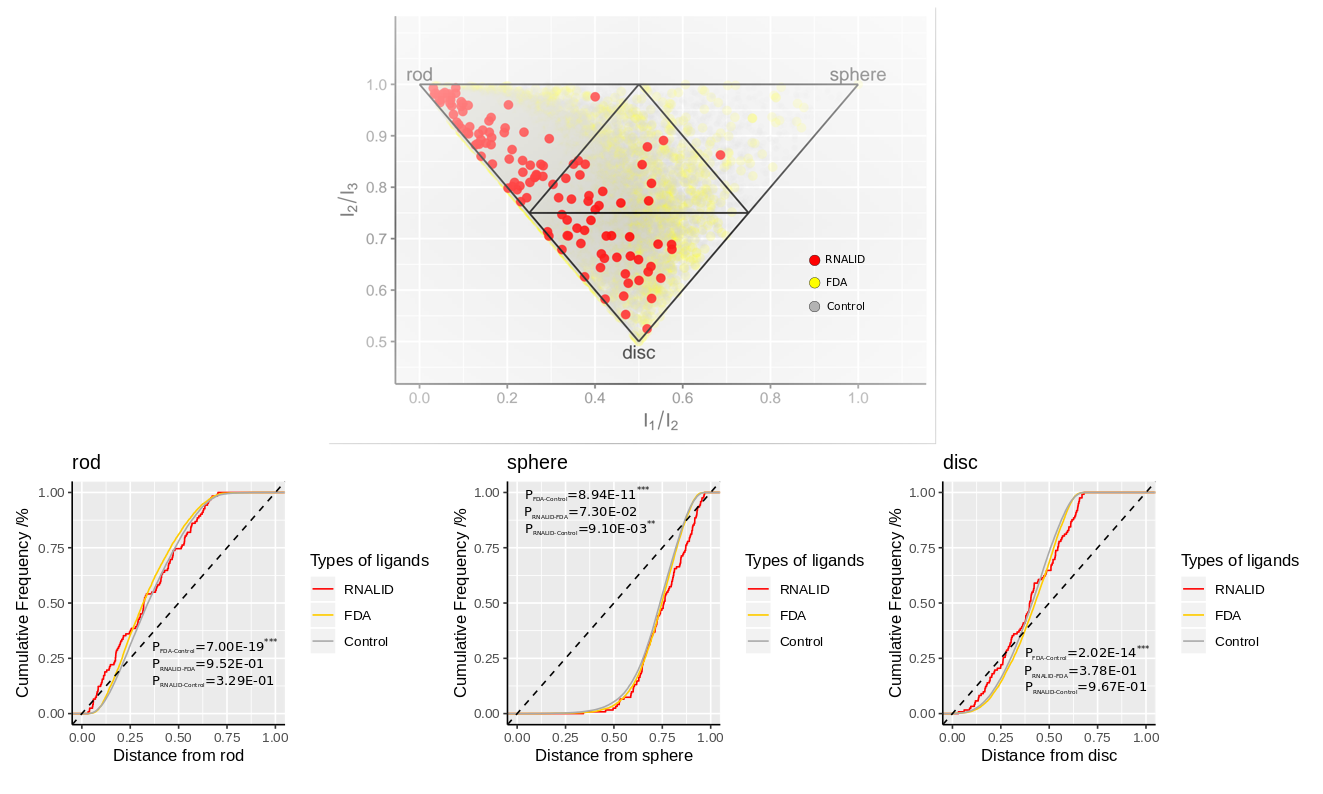

**Figure S10.** PMI analysis comparing ligands in RNALID with FDA approved drugs (protein-binding drugs) and ligands with no bioactivity (control). Student’s t-test was used to evaluate the difference.

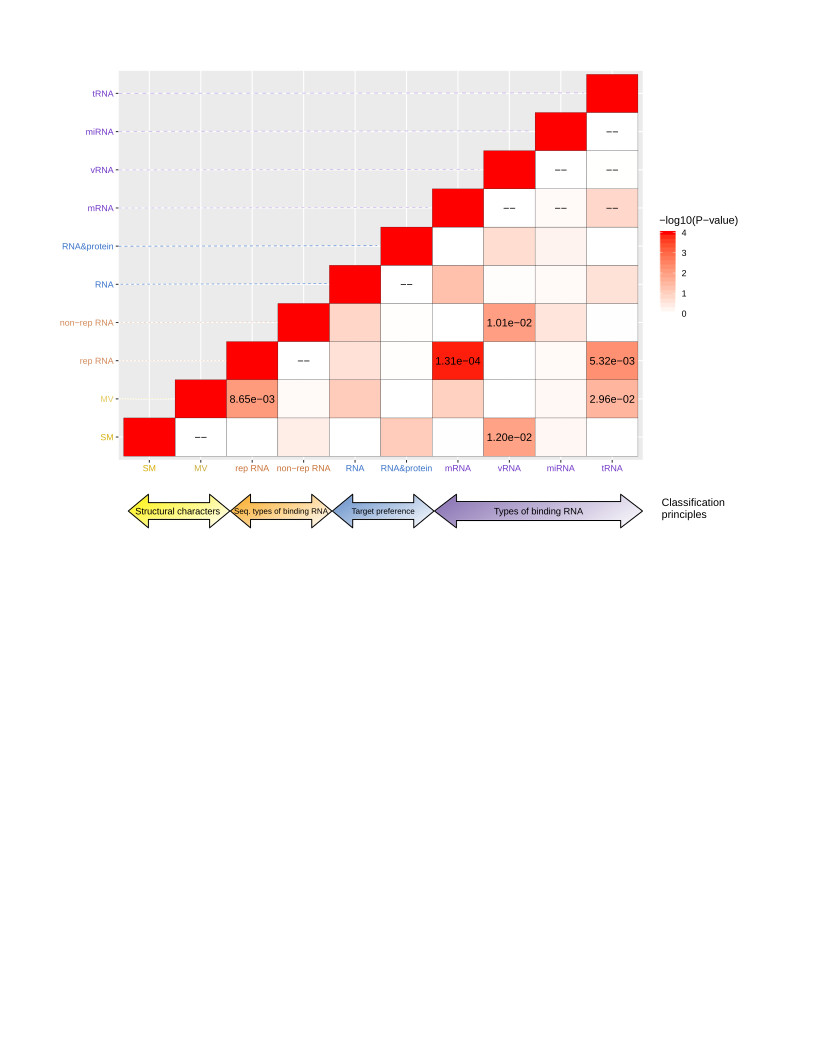

**Figure S11.** Cross ligand-type enrichment analysis. Ligands were classified into different types. “SM” denotes drug-like small molecules, “MV” denotes multivalent ligands, “rep RNA” denotes ligands binding to RNAs containing at least one repeat sequence pattern, “non-rep RNA” denotes ligands binding to RNA not containing repeat sequence, “RNA” denotes ligands only binding with RNA, and “RNA&protein” denotes ligands binding to both RNA and protein. Fisher’s exact test was used to evaluate enrichment of one types of ligands in other types.

**Supplementary tables**

**Table S1.** Sample size of different ligand types. There is overlap among ligand types under the classification principle of “Type of binding RNA”.

| **Classification principles** | **Ligand types** | **Ligand counts** |
| --- | --- | --- |
| Structure characters | SM | 110 |
|  | MV | 18 |
| Target preference | RNA | 104 |
|  | RNA&protein | 24 |
| Sequence types of binding RNA | non-rep RNA | 77 |
|  | rep RNA | 20 |
|  | RNA sequence unavailable | 31 |
| Types of binding RNA | mRNA | 59 |
|  | vRNA | 53 |
|  | tRNA | 11 |
|  | miRNA | 11 |
|  | ncRNA | 7 |
|  | siRNA | 3 |
|  | other RNA types | 5 |

**Table S2.** Functions of RDKit used in calculation of cheminformatic parameters

| **Parameter name** | **RDKit function** | **Detail information** |
| --- | --- | --- |
| MW | Descriptors.ExactMolWt() | Molecular Weight |
| logP | Descriptors.MolLogP() | n-Octanol/Water Partition Coefficient |
| tPSA | Descriptors.TPSA() | Topological Polar Surface Area |
| Fsp3 | rdMolDescriptors.CalcFractionCSP3() | Fraction of sp^3^ Hybridized Carbons |
| ArRings | rdMolDescriptors.CalcNumAromaticCarbocycles() | Number of Aromatic Rings |
| HetRings | rdMolDescriptors.CalcNumAromaticHeterocycles() | Number of Heteroatom-containing Rings |
| nStereo | rdMolDescriptors.CalcNumAtomStereoCenters() | Number of Stereocenters |
| HBA | rdMolDescriptors.CalcNumHBA() | Number of Hydrogen Bond Acceptors |
| HBD | rdMolDescriptors.CalcNumHBD() | Number of Hydrogen Bond Donors |
| Rings | rdMolDescriptors.CalcNumRings() | Number of Rings |
| RotB | rdMolDescriptors.CalcNumRotatableBonds() | Number of Rotatable Bonds |
| ASA | Descriptors.LabuteASA() | Accessible Surface Area |

**Table S3.** Drug-like indexes used in this study

| **Drug-like index** | **Detail** | **Toxicity or unstable drug effect**^c^ |
| --- | --- | --- |
| NR^a^-AR | Androgen receptor | + |
| NR-AR-LBD | Androgen receptor ligand binding domain | + |
| NR-AhR | Aryl hydrocarbon receptor | + |
| NR-Aromatase | CYP19 enzyme | + |
| NR-ER | Estrogen receptors | + |
| NR-ER-LBD | Estrogen receptor alpha ligand-binding domain | + |
| NR-PPAR-gamma | Peroxisome proliferator-activated receptor gamma | + |
| SR^b^-ARE | Anti-oxidative response element | + |
| SR-ATAD5 | ATPase Family AAA Domain Containing 5 | + |
| SR-HSE | Heat shock response element | + |
| SR-MMP | Matrix metalloproteinase | + |
| SR-p53 | P53 | + |
| LOGP | logP | // |
| SOLUBILITY | Solubility | // |
| HERG | Human ether-a-go-go related gene | + |
| LIVER | Liver | + |
| CACO2 | Human Caco-2 cancer cell line | - |
| CYP1A2_I | Cytochrome | + |
| CYP2C9_I | Cytochrome | + |
| CYP2C9_S | Cytochrome | + |
| CYP2C19_I | Cytochrome | + |
| CYP2D6_I | Cytochrome | + |
| CYP2D6_S | Cytochrome | + |
| CYP3A4_I | Cytochrome | + |
| CYP3A4_S | Cytochrome | + |
| BBB | Blood-brain barrier | - |
| CLINTOX | Clinical toxicity | + |
| BACE | Beta-Secretase | + |

^a.^ NR represents nuclear receptors in Tox21 challenge, 2014 (https://tripod.nih.gov/tox21/challenge/)

^b.^ SR represents stress response in Tox21 challenge, 2014 (https://tripod.nih.gov/tox21/challenge/)

^c.^ “+” indicates that QED value is positively related to toxicity or unstable drug effect, “-” indicates a negative relationship, “//” indicates that toxicity or unstable effect would be caused only in a certain range of QED values.

**Table S4.** Drug therapy information of ligands in RNALID.

| **Disease** | **RNA number** | **Drug number** |
| --- | --- | --- |
| Anti-tumoral activities | 1 | 1 |
| Antipruritic; appetite stimulant; antiallergic; post-gastrectomy dumping syndrome | 1 | 1 |
| Bacterial infection | 3 | 3 |
| Cancer | 13 | 11 |
| CHPV infection | 1 | 1 |
| Estrogen agonist or antagonist | 1 | 1 |
| HIV | 1 | 1 |
| Lung disease | 1 | 1 |
| Myotonic dystrophy type 1 | 1 | 1 |
| Parasitic and fungal infections; antidiarrheal | 3 | 2 |
| Pyoderma | 1 | 1 |
| Anthelmintic | 7 | 1 |
| Antiprotozoal agent | 2 | 1 |
| Enhances cAMP-mediated responses in human neuroblastoma cells | 1 | 1 |
| Severe nausea and vomiting; short-term management of psychotic disorders | 2 | 1 |
| Type 1 Spinal Muscular Atrophy (SMA) | 3 | 2 |
| Wound dressings and mouth infections | 1 | 1 |
| Wound dressings; a topical antiseptic | 2 | 1 |

**Table S5.** Details of disease information targeted by ligands in RNALID

| **Disease category** | **Detail** |
| --- | --- |
| Brain or neuro-disease | Alzheimer’s disease |
|  | Parkinson’s disease |
|  | Huntington’s disease |
|  | myotonic dystrophy type 1 (DM1) |
|  | myotonic dystrophy type 2 (DM2) |
|  | fragile X-associated tremor ataxia syndrome (FXTAS) |
|  | Spinal muscular atrophy (SMA) |
| Diabetes and obesity | Diabetes |
|  | Obesity |
| Cancer and tumor | Cancer types |
|  | Tumor types |
| Endometriosis | Endometriosis |
| HCV | Hepatitis C virus |
| HIV | human immunodeficiency virus |
| Microbial | Bacterial or virus inflection other than HCV or HIV |
| Other disease | No acclaim in the publishing |

**Table S6.** Comparison of cheminformatic parameters for FDA approved drugs, ligands without bioactive and ligands in RNALID

| **Chemical characters** | **RNALID vs. FDA** | **RNALID vs. Control** | **FDA vs. Control** |
| --- | --- | --- | --- |
| ArRings | 4.04e-09*** | 3.36e-06*** | 1.28e-15*** |
| ASA | 7.47e-20*** | 4.51e-29*** | 4.65e-81*** |
| Fsp3 | 2.84e-03** | 0.973 | 2.46e-76*** |
| HBA | 8.22e-06*** | 3.55e-12*** | 6.45e-113*** |
| HBD | 2.42e-14*** | 4.54e-38*** | <1.00e-260*** |
| HetRings | 1.33e-11*** | 4.30e-22*** | 2.72e-72*** |
| Logp | 0.114 | 2.01e-04*** | 2.10e-61*** |
| Mw | 6.71e-17*** | 1.15e-26*** | 8.96e-93*** |
| nStereo | 7.28e-03** | 0.347 | 3.90e-194*** |
| Rings | 9.03e-20*** | 3.27e-25*** | 2.59e-17*** |
| RotB | 1.09e-05*** | 2.94e-14*** | 9.56e-171*** |
| tPSA | 1.10e-12*** | 1.21e-26*** | <1.00e-260*** |

MW-U test was used to evaluate the difference. “*” denotes P-value < 0.05, “**” denotes P-value < 0.01, “***” denotes P-value < 0.001.

**Table S7.** Comparison of drug-like indexes for FDA approved drugs, ligands without bioactive and ligands in RNALID.

| **Drug-like indexes** | **RNALID vs. FDA** | **RNALID vs. Control** | **FDA vs. Control** | |
| --- | --- | --- | --- | --- |
| BACE | 1.96e-04*** | 6.07e-04*** | | 0.0557 |
| BBB | 0.243 | 1.49e-06*** | | 2.57e-136*** |
| CACO2 | 0.874 | 1.24e-04*** | | 2.84e-143*** |
| CLINTOX | 1.68e-14*** | 2.62e-12*** | | 1.23e-41*** |
| CYP1A2_I | 0.249 | 3.14e-02* | | 1.88e-125*** |
| CYP2C19_I | 2.73e-04*** | 1.74e-02* | | <1.00e-260*** |
| CYP2C9_I | 9.69e-09*** | 7.61e-14*** | | 6.00e-36*** |
| CYP2C9_S | 1.23e-03** | 1.13e-11*** | | 8.60e-88*** |
| CYP2D6_I | 0.241 | 0.0743 | | 7.15e-04*** |
| CYP2D6_S | 2.13e-05*** | 1.34e-03** | | 1.75e-22*** |
| CYP3A4_I | 3.49e-02* | 0.444 | | 6.91e-22*** |
| CYP3A4_S | 2.85e-02* | 1.35e-02* | | 0.14 |
| HERG | 9.28e-11*** | 0.183 | | <1.00e-260*** |
| LIVER | 0.121 | 0.439 | | 1.09e-67*** |
| LOGP | 3.22e-05*** | 1.07e-21*** | | <1.00e-260*** |
| NR.AhR | 6.67e-17*** | 9.64e-11*** | | 1.37e-87*** |
| NR.AR | 3.78e-35*** | 3.29e-46*** | | 6.30e-159*** |
| NR.AR.LBD | 2.03e-19*** | 1.51e-22*** | | 1.27e-09*** |
| NR.Aromatase | 2.17e-26*** | 4.81e-21*** | | 2.93e-70*** |
| NR.ER | 1.49e-07*** | 1.20e-03** | | 6.97e-42*** |
| NR.ER.LBD | 5.58e-21*** | 1.39e-14*** | | 2.57e-39*** |
| NR.PPAR.gamma | 1.16e-12*** | 2.22e-06*** | | 6.71e-105*** |
| SOLUBILITY | 4.41e-03** | 1.33e-03** | | <1.00e-260*** |
| SR.ARE | 2.20e-20*** | 1.05e-12*** | | 3.38e-91*** |
| SR.ATAD5 | 1.22e-21*** | 1.59e-13*** | | 9.10e-102*** |
| SR.HSE | 2.21e-35*** | 2.51e-16*** | | <1.00e-260*** |
| SR.MMP | 2.20e-14*** | 1.24e-05*** | | 1.97e-165*** |
| SR.p53 | 3.35e-11*** | 1.04e-08*** | | 4.71e-15*** |

MW-U test was used to evaluate the difference. “*” denotes P-value < 0.05, “**” denotes P-value < 0.01, “***” denotes P-value < 0.001.

**Table S8.** Information of ligands with novel structures in RNALID.

| **Ligand Name** | **RNA name** | **RNA type** | **Disease** | **Ligand types** | **Reference(PMID)** |
| --- | --- | --- | --- | --- | --- |
| ZINC04574788 | miR-21 pre-element | miRNA | Endometriosis | SM, RNA, non-rep RNA | 29625267 |
| Regorafenib | miR-21 pre-element | miRNA | cancer | SM, RNA&protein, non-rep RNA | 28236743 |
| 5-(N,N)-Dimethyl amiloride (6) | HIV-1 TAR RNA | vRNA | HIV | SM, RNA, non-rep RNA | 21706033 |
| cationic porphyrin (5,10,15,20-tetra(N-methyl-4-pyridyl) porphyrin (TMPyP4)) | MT3-MMP<U+00A0>mRNA （any RNA with r(GGGGCC)n that forms a quadruplex structure） | mRNA, siRNA | neurological disease | SM, RNA | 24371143 |
| Negamycin (NEG) （2-(3,6-diamino-5-hydroxyhexanoyl)-methylhydrazino]acetic acid） | E. coli<U+00A0>tRNA | tRNA | bacteria | SM, RNA | 25306922 |
| dinuclear iron (II) metallosupramolecular triple helicates [Fe2L3]Cl4<U+00A0>(L<U+2009>=<U+2009>C25H20N4) | HIV-1 TAR RNA | vRNA | HIV | SM, RNA, non-rep RNA | 27405089 |
| 4,5-trisubstituted oxazolidinones | AM1A | ncRNA | bacteria | SM, RNA, non-rep RNA | 21812425 |
| Azidothymidine | Yeast Baker RNA | yeast RNA | HIV | SM, RNA&protein | 15585137 |
| Antofine | TMV RNA | vRNA | Virus | SM, RNA, non-rep RNA | 25154018, 22753104,  25809229 |
| X2SS | pre-miRNA | miRNA | -- | SM, RNA | 23164709 |
| ARGININE | HIV-1 TAR RNA | vRNA | HIV | SM, RNA, non-rep RNA | 21574948 |
| ATF_compound12 | TMV RNA | vRNA | Virus | SM, RNA&protein, rep RNA | 25809229 |

**References**

1.	Morgan, B. S., et al., R-BIND: An Interactive Database for Exploring and Developing RNA-Targeted Chemical Probes. *ACS Chem Biol* **2019,** *14* (12), 2691-2700.

2.	Morgan, B. S., et al., Discovery of Key Physicochemical, Structural, and Spatial Properties of RNA-Targeted Bioactive Ligands. *Angew Chem Int Ed Engl* **2017,** *56* (43), 13498-13502.

3.	Lorenz, R., et al., ViennaRNA Package 2.0. *Algorithms Mol Biol* **2011,** *6*, 26.

4.	Camacho, C., et al., BLAST+: architecture and applications. *BMC Bioinformatics* **2009,** *10*, 421.

5.	Knox, C., et al., DrugBank 3.0: a comprehensive resource for 'omics' research on drugs. *Nucleic Acids Res* **2011,** *39* (Database issue), D1035-41.

6.	Law, V., et al., DrugBank 4.0: shedding new light on drug metabolism. *Nucleic Acids Res* **2014,** *42* (Database issue), D1091-7.

7.	Wishart, D. S., et al., DrugBank 5.0: a major update to the DrugBank database for 2018. *Nucleic Acids Res* **2018,** *46* (D1), D1074-D1082.

8.	Wishart, D. S., et al., DrugBank: a knowledgebase for drugs, drug actions and drug targets. *Nucleic Acids Res* **2008,** *36* (Database issue), D901-6.

9.	Wishart, D. S., et al., DrugBank: a comprehensive resource for in silico drug discovery and exploration. *Nucleic Acids Res* **2006,** *34* (Database issue), D668-72.

10.	Yang, Y., et al., SPOT-Ligand: Fast and effective structure-based virtual screening by binding homology search according to ligand and receptor similarity. *J Comput Chem* **2016,** *37* (18), 1734-9.

11.	Fan, C., et al., DStruBTarget: Integrating Binding Affinity with Structure Similarity for Ligand-Binding Protein Prediction. *J Chem Inf Model* **2020,** *60* (1), 400-409.

12.	Hu, J., et al., LS-align: an atom-level, flexible ligand structural alignment algorithm for high-throughput virtual screening. *Bioinformatics* **2018,** *34* (13), 2209-2218.
